# Mild traumatic brain injury induces transient, sequential increases in proliferation, neuroblasts/immature neurons, and cell survival: a time course study in the male mouse dentate gyrus

**DOI:** 10.1101/2020.10.07.330118

**Authors:** LR Clark, S Yun, NK Acquah, PL Kumar, HE Metheny, RCC Paixao, AS Cohen, AJ Eisch

## Abstract

Mild traumatic brain injuries (mTBIs) are prevalent worldwide. mTBIs can impair hippocampal-based functions such as memory and cause network hyperexcitability of the dentate gyrus (DG), a key entry point to hippocampal circuitry. One candidate for mediating mTBI-induced hippocampal cognitive and physiological dysfunction is injury-induced changes in the process of DG neurogenesis. There are conflicting results on how TBI impacts the process of DG neurogenesis; this is not surprising given that both the neurogenesis process and the post-injury period are dynamic, and that the quantification of neurogenesis varies widely in the literature. Even within the minority of TBI studies focusing specifically on mild injuries, there is disagreement about if and how mTBI changes the process of DG neurogenesis. Here we utilized a clinically-relevant rodent model of mTBI (lateral fluid percussion injury, LFPI), gold-standard markers and quantification of the neurogenesis process, and three time points post-injury to generate a comprehensive picture of how mTBI affects adult hippocampal DG neurogenesis. Male C57BL/6J mice (6-8 weeks old) received either sham surgery or mTBI via LFPI. Proliferating cells, neuroblasts/immature neurons, and surviving cells were quantified via stereology in DG subregions (subgranular zone [SGZ], outer granule cell layer [oGCL], molecular layer, and hilus) at short-term (3 days post-injury, dpi), intermediate (7 dpi), and long-term (31 dpi) time points. The data suggest this model of mTBI induces transient, sequential increases in ipsilateral SGZ/GCL proliferating cells, neuroblasts/immature neurons, and surviving cells which are suggestive of mTBI-induced neurogenesis. In contrast to these ipsilateral hemisphere findings, measures in the contralateral hemisphere were not increased in key neurogenic DG subregions after LFPI. Our work in this mTBI model is in line with most literature on other and more severe models of TBI in showing TBI stimulates the process of DG neurogenesis. However, as our DG data in mTBI provide temporal, subregional, and neurogenesis-stage resolution, these data are important to consider in regard to the functional importance of TBI-induction of the neurogenesis process and future work assessing the potential of replacing and/or repairing DG neurons in the brain after TBI.

## 1. INTRODUCTION

Traumatic brain injury (TBI) is a leading cause of death and disability in the United States, affecting ~2.8 million new people annually (Report to Congress on Mild Traumatic Brain Injury in the United States; Taylor et al., 2017). TBI is a mechanical event that results in primary focal and/or diffuse brain injury, often with secondary pathological sequelae and long-term neurological consequences (Sterr et al., 2006; Daneshvar et al., 2011; Katz et al., 2015). TBIs in humans are heterogeneous in location, severity, and mechanism, but are typically classified as mild, moderate, or severe based on severity and duration of acute symptoms (Narayan et al., 2002). Most research has focused on moderate and severe injuries due to their dramatic impact on brain health and cognition (Lippert-Grüner et al., 2006). However, the majority of brain injuries sustained are classified as mild TBIs (mTBIs)(Faul et al., 2010; Taylor et al., 2017). mTBIs induce no gross structural brain abnormalities, but can lead to lasting cognitive deficits - particularly in regard to attention and memory (Daneshvar et al., 2011; Katz et al., 2015). Injury-induced cognitive deficits in the absence of gross brain damage can be replicated in rodent models of mTBI (Lyeth et al., 1990; Eakin and Miller, 2012). Notably, many of the cognitive deficits seen in both rodents and humans after TBI involve hippocampal-dependent functions, such as spatial and contextual memory and pattern separation (Witgen et al., 2005; Smith et al., 2012; Folweiler et al., 2018; Paterno et al., 2018). These cognitive deficits are accompanied by electrophysiological changes in the hippocampus, including hyperexcitability in the dentate gyrus (DG; Folweiler et al., 2018). The DG typically acts as a “filter” or “gate”, with sparse activity limiting how much information is relayed through to downstream areas (Hsu, 2007). One theory about mTBI-induced cognitive deficits posits that this “DG gate” breaks down after injury, leading to excessive DG depolarization which spreads to area CA3 (Folweiler et al., 2018), thus disrupting this “gating” of information flow through the hippocampus and causing cognitive dysfunction. Although an excitatory/inhibitory imbalance is thought to mediate injury-induced cognitive deficits, the cellular and network mechanisms underlying these mTBI-induced physiological and cognitive deficits are unknown.

One potential contributor to rodent mTBI-induced physiological changes and cognitive deficits is mTBI-induced changes in the generation of new hippocampal DG granule cells. This process of DG neurogenesis consists of ‘stages’, including proliferation (cell division of stem cells and their progeny in the subgranular zone [SGZ]), generation of neuroblasts and immature neurons, and cell survival of mature, glutamatergic DG granule cells (Aimone et al., 2014; Bond et al., 2015; Kempermann et al., 2015). The bidirectional relationship between the activity of the DG as a whole and the process of neurogenesis has been well-studied (Ma et al., 2009; Ikrar et al., 2013; Kempermann, 2015; Temprana et al., 2015). While the exact function of adult-born DG granule cells is under debate, they are implicated in hippocampal-based cognition such as contextual memory and pattern separation (Clelland et al., 2009; Tronel et al., 2010; Sahay et al., 2011; Nakashiba et al., 2012). The process of DG neurogenesis also contributes to circuit excitability in both physiological (Ikrar et al., 2013) and pathological conditions like epilepsy (Cho et al., 2015; Neuberger et al., 2017). Given that the process of neurogenesis has potent influence over hippocampal physiology and function, there is great interest in understanding how it - and associated hippocampal physiology and function - is changed after brain injury.

Indeed, many preclinical studies suggest the process of adult hippocampal DG neurogenesis is changed after TBI (Aertker et al., 2016; Ngwenya and Danzer, 2018; Bielefeld et al., 2019). However, the direction of the change is disputed. TBI is reported to decrease (Gao et al., 2008; Hood et al., 2018), increase (Dash et al., 2001; Kernie et al., 2001; Blaiss et al., 2011; Villasana et al., 2015; Sun, 2016; Neuberger et al., 2017), or not change (but rather increases in glial cell proliferation; Chirumamilla et al., 2002; Rola et al., 2006) the process of DG neurogenesis. These discrepancies are likely related in part to differences in experimental parameters, including time point post-TBI, ‘stage’ of the process examined, approach to quantify the neurogenesis process, and TBI model used and its severity. To this end, it is notable that rodent models of mild brain injury - including lateral fluid percussion injury (LFPI), blast trauma, and weight drop - also produce mixed effects on the process of DG neurogenesis (Bye et al., 2011; Wang et al., 2016; Neuberger et al., 2017; Tomura et al., 2020). Given that new neuron replacement has been suggested as a potential treatment for TBI-induced cognitive dysfunction (Blaiss et al., 2011; Sun et al., 2015; Aertker et al., 2016; Sun, 2016), and the prevalence of mTBI in humans, it is surprising that only two rodent studies have examined injury-induced effects on the process of DG neurogenesis after mTBI caused by LFPI. One LFPI study in mice reported the number of neuroblasts/immature neurons labeled by doublecortin (DCX) in the ipsilateral DG increased to the same magnitude in Sham and mTBI mice vs. control mice (Aleem et al., 2020). However, this study only examined DCX-immunoreactive (+) cells 7 days post-injury (dpi), which urges study of additional time points; also this study did not provide neurogenesis quantification methods, making it unclear whether this neurogenesis result is meaningful. A very comprehensive LFPI study in rat sampled 4 time points post-injury and used stereology to quantify cells labeled with markers of several stages of the neurogenesis process, and definitely showed a transient increase in DCX+ cells 3 dpi (Neuberger et al., 2017). However, this study was performed in juvenile rats, leaving open the question of how mTBI changes the process of neurogenesis in the adult mouse. In light of the cognitive and physiological hippocampal changes seen in the mTBI mouse model (Witgen et al., 2005; Smith et al., 2012; Folweiler et al., 2018; Paterno et al., 2018), it is important to address this major knowledge gap: how does a clinically-relevant model of mTBI change the dynamic process of DG neurogenesis in the adult mouse?

To fill this knowledge gap, we exposed adult male mice to LFPI or Sham and examined stages of DG neurogenesis at short-term (3 dpi), intermediate (7 dpi), and long-term (31 dpi) time points. The clinically-relevant model LFPI was used because it causes two changes in mice that are also seen in humans after mTBI: 1) both focal and diffuse injury but no necrotic cavity (Smith et al., 2012; Xiong et al., 2013; Brady et al., 2018) and 2) cognitive deficits, such as worse hippocampal-based spatial memory (Folweiler et al., 2018). In our study, we collected indices of proliferation (number of Ki67+ cells at all time points, number of BrdU+ cells 3 dpi), neuroblasts/immature neurons (number of DCX+ cells at all time points), and cell survival (number of BrdU+ cells 7 and 31 dpi). We find the ipsilateral DG neurogenic regions of LFPI mice have more proliferation 3 dpi, more neuroblasts/immature neurons 7 dpi, and more new cell survival 31 dpi relative to these regions in Sham mice. These results suggest this model of mTBI produces a transient increase of DG neurogenesis in canonical DG neurogenic regions, consistent with the increased neurogenesis seen after other and more severe models of TBI. We discuss this and other interpretations of our DG data, and consider the implications for the temporal, subregional, and neurogenesis-stage resolution data provided here for the first time in a mouse model of mTBI.

## 2. METHODS

### 2.1 Mice

Experiments were performed on 6- to 8-week-old male C57BL/6J mice (Jackson Laboratory, Bar Harbor, ME; IMSR JAX:000664, RRID:IMSR_JAX:000664, **Fig. 1A**). Mice were group-housed 5/cage in an AAALAC-approved facility at Children’s Hospital of Philadelphia (CHOP). The vivariums at the Colket Translational Research Building (Penn/CHOP) are temperature- and humidity-controlled. Lights are on at 6am and off at 6pm, and food and water are provided *ad libitum*. All experiments were carried out in accordance with protocols approved by the Institutional Animal Care and Use Committee of CHOP and the guidelines established by the NIH Guide for the Care and Use of Laboratory Animals.

**Figure 1.**
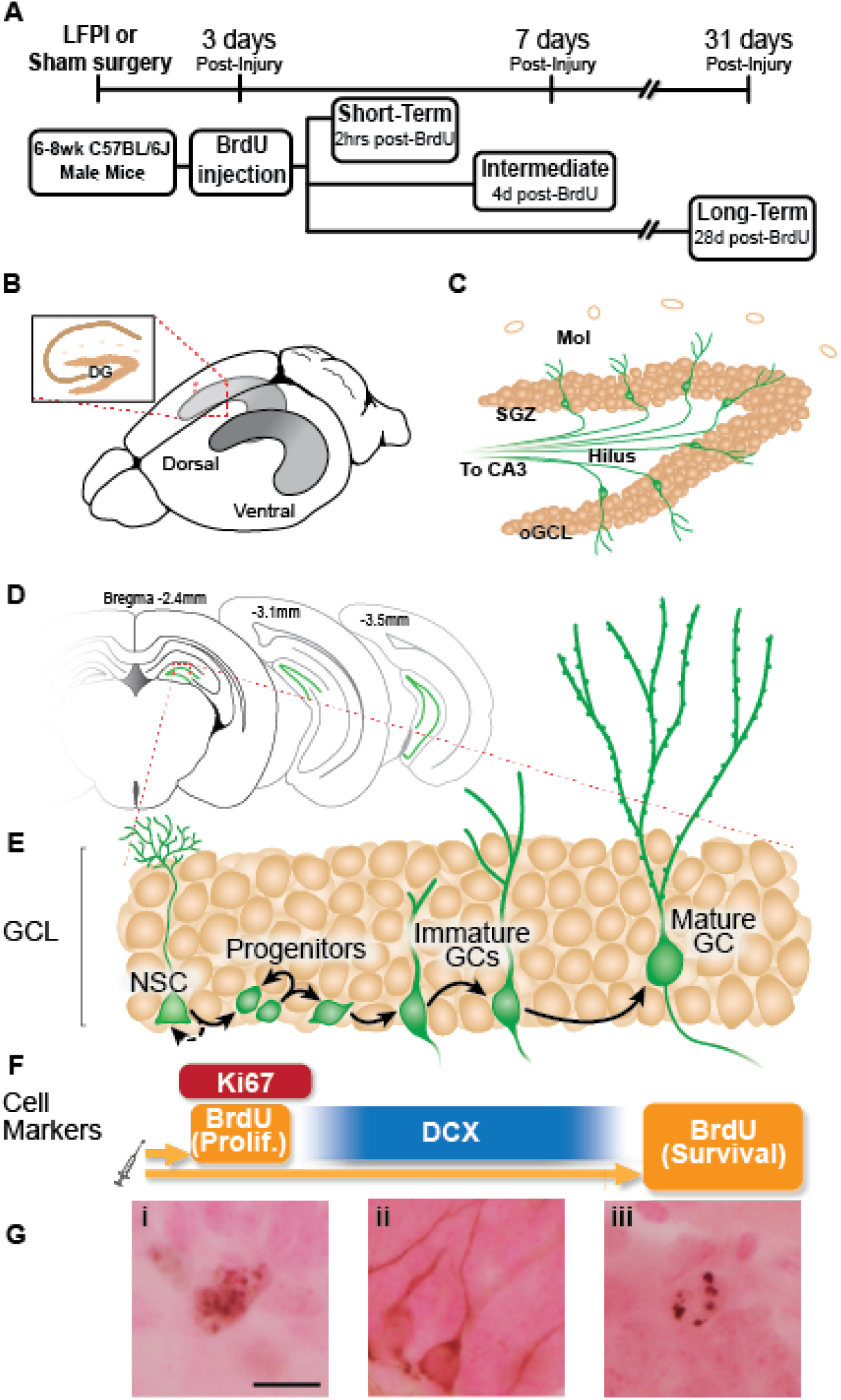
Experimental timeline and overview of dentate gyrus (DG) neurogenesis and immunohistochemical markers used. **(A)** Timeline of experimental procedures. 6-8 wk old male C57BL6/J mice received lateral fluid percussive injury (LFPI) or Sham surgery, and 150 mg/kg i.p. Bromodeoxyuridine (BrdU) injection 3 days post-injury (dpi). The Short-Term group was perfused 3 dpi (2 hours [hrs] post-BrdU); the Intermediate group perfused 7 dpi (4 days post-BrdU); and the Long-Term group perfused 31 dpi (28 days post-BrdU). **(B)** Schematic of a whole mouse brain showing bilateral location of the hippocampus (grey structure). Exploded inset (red dotted lines) depicts a coronal section through the dorsal hippocampus (tan indicates primary cell layers) with the granule cell layer (GCL) of the DG indicated. **(C)** Schematic of enlarged DG subregions. GCL is represented by densely-packed tan circles; individual tan circles represent dorsal boundary of DG (hippocampal fissure). Green circles/lines are GCL GCs and their axons project through the Hilus to eventually reach CA3 (not pictured). Mol: molecular layer. SGZ: subgranular zone. oGCL: outer granule cell layer. **(D)** Schematic of location of DG (green lines) in coronal sections of a mouse brain at three distances from Bregma (−2.4, −3.1, −3.5 mm). **(E-F)** Exploded inset of red dotted line in **(D)** depicts stages of GCL neurogenesis and the antibodies with which cells in each neurogenesis stage can be labeled. Neural stem cells (NSCs) asymmetrically divide to create progenitor cells, whose progeny differentiate into immature granule cells, some of which survive to become mature granule cells. BrdU (in the Short-Term group) and Ki67 (all groups) mark dividing cells; doublecortin (DCX) marks neuroblasts/immature neurons (in all groups); and BrdU in the Long-term group marks surviving cells (cells that were dividing and labelled at the time of BrdU injection, and have survived to 31 dpi). **(G)** Representative photomicrographs of cells stained with antibodies against Ki67 **(Gi)**, DCX **(Gii)**, or BrdU **(Giii)**. Scale bar = 10um.

### 2.2 Craniectomy

All mice underwent craniectomy on day −1 of the experiment (Dixon et al., 1987; Smith et al., 2012). Mice were anesthetized with an intraperitoneal (i.p.) injection of ketamine (100 mg/kg) and xylazine (6-16 mg/kg). Once anesthetized, mice were placed in a stereotaxic frame (Stoelting, Wood Dale, IL, USA), the scalp was incised and pulled away to fully expose the right parietal bone. An ultra-thin Teflon disk (3-mm outer diameter) was positioned between Lambda and Bregma and between the sagittal suture and the lateral ridge over the right hemisphere, and then glued to the skull with Vetbond (3M, St. Paul, MN, USA). Guided by the Teflon disk, a trephine was used to perform a 3-mm diameter craniectomy over the right parietal area. Following craniectomy, a Luer-lock needle hub (3-mm inner diameter) was secured above the skull opening with Loctite superglue and dental acrylic, filled with saline and capped. Mice were removed from the stereotaxic apparatus, placed on a heating pad until fully recovered from anesthesia, and then returned to their respective home cage.

### 2.3 Lateral fluid percussion injury (LFPI)

Twenty-four hours (hrs) post-craniectomy, mice underwent LFPI or Sham surgery (Fig. 1A; Dixon et al., 1987; Smith et al., 2012). Mice were placed under isoflurane anesthesia (2% oxygen, 500ml/min) in a chamber. Respiration was visually monitored until mice reached a surgical plane of anesthesia (1 respiration/2 sec). Mice were then removed from isoflurane and the needle hub was refilled with saline and connected to the fluid percussion injury device (Department of Biomedical Engineering, Virginia Commonwealth University) via high-pressure tubing. The mouse was placed onto a heating pad on its left side. On resumption of normal breathing pattern but before sensitivity to stimulation, the injury was induced by a 20-msec pulse of saline onto the intact dura. The pressure transduced onto the dura was monitored with an oscilloscope, with injury severity ranging from 1.4 to 1.6 atmospheres. Sham mice underwent all surgical procedures including attachment to the FPI device with exclusion of the actual fluid pulse. Immediately after injury or sham surgery, the hub was removed from the skull and the mouse was placed in a supine position to measure the latency to righting. All mice in this study righted themselves within 3 min, while Sham mice righted themselves in <1 min. After righting, the mouse was returned to placement under isoflurane for scalp suturing. Mice recovered on a heating pad until mobile, at which point they were returned to their home cage. Within a cage, mice were exposed to either Sham or LFPI and post-surgery Sham and LFPI mice were housed in the same cage. Two cohorts of 5 Sham and 5 LFPI mice were generated for each condition at each time point (3, 7, and 31 days post-Sham or post-LFPI).

### 2.4 Bromodeoxyuridine (BrdU) administration

BrdU (Accurate Scientific, OBT0030G), a thymidine analog, was freshly-prepared at 10mg/mL in 0.09% sterile saline and 0.007N NaOH. All mice received a single 150 mg/kg i.p. injection 3 days after Sham or LFPI **(Fig. 1A)**. Mice were weighed the morning of BrdU administration, and left undisturbed until BrdU injection was delivered that afternoon, and also were undisturbed for 2 hours post-BrdU injection. In C57BL/6J male mice, this BrdU dose and injection procedure ‘pulse-labels’ cells in S-phase of the cell cycle at the time of BrdU injection (Mandyam et al., 2007).

### 2.5 Tissue collection and immunohistochemistry (IHC)

Three, 7, or 31 days post-Sham or -LFPI **(Fig. 1)**, mice were anesthetized with chloral hydrate (400 mg/kg, i.p.) prior to intracardial perfusion with ice-cold 0.1 M phosphate-buffered saline (PBS) for exsanguination followed by 4% paraformaldehyde for fixation (Rivera et al., 2013; DeCarolis et al., 2014). Extracted brains were immersed for 24-hrs in 4% paraformaldehyde in 0.1M PBS at 4°C for post-fixation, followed by least 3 days of immersion in 30% sucrose in 0.1 M PBS for cryoprotection with 0.01% sodium azide to prevent bacterial growth. For coronal brain sectioning, the brain of each mouse extending from anterior to the DG to the cerebellum (from 0.22 to −5.34 μm from Bregma) was sectioned at 30 μm in a 1:9 series using a freezing microtome (Leica SM 2000 R Sliding Microtome). Sections were stored in 1xPBS with 0.01% sodium azide at 4°C until processing for IHC. Slide-mounted IHC for Ki67+, DCX+, and BrdU+ cells in the DG was performed as previously described (Rivera et al., 2013; DeCarolis et al., 2014). Briefly, one entire series of the hippocampus (every 9th section) was slide-mounted onto charged slides (Fisher Scientific, 12-550-15, Pittsburgh, PA). Slide-mounted sections underwent antigen retrieval (0.01 M citric acid pH 6.9, 100°C, 15 min) followed by washing in 1xPBS at room temperature. For BrdU IHC, two additional steps were performed to allow the antibody access to DNA inside the cell nucleus: permeabilization (0.1% Trypsin in 0.1 M TRIS and 0.1% CaCl2, 10 min) and denaturation (2N HCl in 1x PBS, 30 min). Non-specific binding was blocked with 3% serum (donkey) and 0.3% Triton-X in PBS for 30 min. After blocking and pretreatment steps, sections were incubated with rat-α-BrdU (1:400; Accurate catalog OBT0030, Westbury, NY), rabbit-α-Ki67 antibody (1:500; Fisher Scientific catalog RM-9106S, Freemont, CA), or goat-α-DCX (1:4000; Santa Cruz Biotechnology catalog sc-8066, Dallas, TX) in 3% serum and 0.3% Tween-20 overnight. For single labeling IHC, primary antibody incubation was followed by 1xPBS rinses, incubation with biotinylated secondary antibodies (biotin-donkey-α-rat-IgG, catalog 712-065-153; biotin-donkey-α-rabbit-IgG, catalog 711-065-152; or biotin-donkey-α-goat-IgG, catalog 705-065-003; all 1:200, all from Jackson ImmunoResearch, West Grove, PA) for 1 hr, and 1xPBS rinses. Next, endogenous peroxidase activity was inhibited via incubation with 0.3% hydrogen peroxide (H_2_O_2_) for 30 min, followed by incubation with an avidin-biotin complex for 60-90 min (ABC Elite, Vector Laboratories PK-6100). After another set of rinses in 1xPBS, immunoreactive cells were visualized via incubation with metal-enhanced diaminobenzidine (Fisher Scientific, 34065, Pittsburgh, PA) for 5-10 min. Finally, slides were incubated for ~2 min in the nuclear counterstain, Fast Red (Vector Laboratories catalog H3403), dehydrated via a series of increasing ethanol concentrations, and coverslipped using DPX (Fisher Scientific, 50-980-370, Pittsburgh, PA).

### 2.6 Cavalieri volume estimation

Granule cell layer (GCL) volume was assessed using the Cavalieri Estimator Probe within the Stereo Investigator software (Gundersen et al., 1988; Harburg et al., 2007; Basler et al., 2017). All measurements were obtained using the Stereo Investigator software (MBF Bioscience, Williston, VT) and a 40x objective (numerical aperture [NA] 0.75) on a Zeiss AxioImager M2 microscope. A 20 μm^2^ counting grid was superimposed over a live image of each section that contained the DG. This grid was used to calculate the area of the GCL at each distance from Bregma. Using the Cavalieri principle (Gundersen and Jensen, 1987; Gundersen et al., 1988), the area values were used to find the volume of the DG GCL.

### 2.7 Cell type quantification

Stereology was used to quantify indices relevant to the process of DG neurogenesis (Ki67+, DCX+, and BrdU+ cell number), as described in each subsection below and shown in **Fig. 1B-G**.

#### 2.7.1 Quantification of Ki67+ and BrdU+ cells in the DG

Due to their rarity (Lagace et al., 2010), BrdU+ and Ki67+ cells were quantified exhaustively in the DG in every 9th section spanning the entire hippocampus via brightfield microscopy using a 40X objective (NA 0.90) on an Olympus BX 51 microscope. Factors considered in determining Ki67+ or BrdU+ cells were size, color, shape, transparency, location, and focal plane. BrdU+ and Ki67+ cells were quantified in 4 DG subregions: the SGZ (40um into the hilus and the inner half of the GCL), widely considered to be the “neurogenic niche” of the DG (Riquelme et al., 2008; Zhao et al., 2008; Kezele et al., 2017; Obernier and Alvarez-Buylla, 2019); the outer GCL (oGCL), to which a minority of adult-generated cells migrate (Kempermann et al., 2003); the hilus, through which DG granule cells project their processes toward CA3; and the molecular layer, the site of DG granule cell dendrites and DG inputs from the entorhinal cortex. Resulting cell counts were multiplied by 9 to get total cell number.

#### 2.7.2 Quantification of DCX+ cells in the DG

DCX+ cells in the DG were quantified via unbiased stereology in only one region of interest (ROI): the SGZ and GCL proper (including the inner and oGCL). DCX+ cells in the oGCL were rare, and thus were considered part of the GCL. DCX+ cells in the hilus and molecular layer had ambiguous/faint staining with this and other antibodies, and thus DCX+ cells in these regions were not quantified. DCX+ cells in the SGZ/GCL are densely presented compared to the relatively rare populations of Ki67+ and BrdU+ cells; therefore, DCX+ cells in the SGZ/GCL were quantified using the optical fractionator workflow in StereoInvestigator (MicroBrightField, MBF) (Brown et al., 2003; Lagace et al., 2010; Zhao et al., 2010). Briefly, every 9th coronal section spanning the entire hippocampus was analyzed on a Nikon Eclipse Ni or Zeiss AxioImager M2 microscope. The ROI (SGZ/GCL) was traced for each section at 100X magnification (10X objective, NA 0.30). Quantification was performed at 400X (40X objective, NA 0.75) by focusing through the Z-plane of the section. DCX+ cells were counted if they fulfilled three criteria: the entire boundary of the cell body was visible, there was a dendritic process emerging from the cell body, and the soma was darker than the surrounding background. DCX+ cells were only excluded on the basis of size if they were small enough to be a swelling of a dendrite (< ~5um).

#### 2.7.3 Quantification of Ki67+, BrdU+, and DCX+ cells along the longitudinal axis (by Bregma)

For anterior/posterior analysis of Ki67+, BrdU+, and DCX+ cell number, results were divided into anterior and posterior bins, defined by a division at Bregma level −2.60mm (Tanti et al., 2012; Tanti and Belzung, 2013; Zhou et al., 2016). Bregma level −2.60mm was identified based on when hippocampal CA3 reaches approximately halfway down the dorsal/ventral extent of the brain, and the corpus callosum no longer connects the left and right hemispheres (Paxinos and Franklin, 2019). Cell counts for each section anterior to −2.60mm were summed and multiplied by 9 to get an anterior value for each mouse. The same procedure was used for the posterior analysis, using all sections posterior to Bregma level −2.60mm.

### 2.8 Data presentation and analysis and image presentation

Experimenters were blinded to injury condition, and code was only broken after data analyses were complete. Counts for cellular markers were collected for at least 5 (cohort 1) and at most 10 (cohorts 1 and 2) Sham and LFPI mice at the 3, 7, and 31-dpi time points. Due to the pandemic-decreed, multi-month lab shut-down, not every cellular marker at each time point was analyzed in both cohorts. However, for cellular markers that were quantified in both cohorts, the data from each cohort were pooled together after confirmation of no cohort variance. Data are presented as individual data points with mean and standard error of the mean displayed. Data were assessed for normality with the Shapiro-Wilk test. Normal data were analyzed with two-tailed t-test, and non-normal data were analyzed with the Mann-Whitney test. Graphs were generated in GraphPad Prism (version 8.0). Full details of statistical analyses for each figure panel are provided in Tables S1 and S2. Photomicrographs were taken with an Olympus DP74 camera using CellSens Standard software, or a Zeiss Lumina HR color camera using MBF StereoInvestigator software, and imported into Adobe Illustrator (version 24.3) for cropping and labeling. For photomicrographs in Figures 2, 4, and 6, representative images from similar Bregma levels are provided when the difference between Sham and LFPI cell numbers was significant. For Figures 3, 5, and 7, a schematic of the DG with a given subregion outlined in red is included to the left of the corresponding cell counts.

## 3. RESULTS

### 3.1 Weight and gross locomotor activity 3, 7, and 31 dpi

This LFPI mouse model of mild TBI (Dixon et al., 1987; Smith et al., 2012) did not change weight or weight gain between Sham and LFPI mice when at the three time points examined: 3, 7, or 31 dpi (**Fig. 1A**; data not shown). Observation several hours prior to perfusion at each time point by an experimenter blinded to injury condition did not reveal gross differences in locomotor activity between Sham and LFPI mice.

### 3.2 Three dpi: neurogenesis indices in the ipsilateral DG neurogenic regions

Three days was selected as the earliest time point post-Sham or post-LFPI to quantify indices of DG neurogenesis (proliferation and neuroblasts/immature neurons). This was for two reasons. First, DG function and neurogenesis are commonly studied 7 dpi (Gilley and Kernie, 2011; Sun et al., 2015; Ibrahim et al., 2016; Neuberger et al., 2017; Shapiro, 2017; Carlson and Saatman, 2018; Wu et al., 2018; Aleem et al., 2020), a time point when the DG is hyperexcitable and DG-dependent spatial memory is impaired (Folweiler et al., 2018). DG neurogenesis influences both DG excitability and DG-dependent behavior (Lacefield et al., 2012; Ikrar et al., 2013; Park et al., 2015; Neuberger et al., 2017). Therefore, examination of neurogenesis indices 3 dpi may reveal changes that influence the subsequent emergence of 7 dpi DG hyperexcitability and functional impairment. Second, indices of DG proliferation and neurogenesis are changed 3 dpi in more severe TBI models relative to control animals (Dash et al., 2001; Chirumamilla et al., 2002; Peters et al., 2018; Villasana et al., 2019), but only a few papers look at DG neurogenesis at an short-term time point post-mTBI, and the results are mixed (Wang et al., 2016; Neuberger et al., 2017; Tomura et al., 2020). To address this knowledge gap, the neurogenic regions of the DG (SGZ, GCL) from Sham and LFPI mice were assessed 3 dpi for indices of proliferation (Ki67+ and BrdU+ cell number) and neuroblasts/immature neurons (DCX+ cell number).

#### 3.2.1 Three dpi, there are more Ki67+ proliferating cells (cells in the cell cycle) in the ipsilateral neurogenic region (SGZ) in LFPI mice relative to Sham mice

Ki67 is an endogenous protein with a short half-life expressed in all cells that are in stages of the cell cycle (G1, S, G2, M)(Brown and Gatter, 2002). Ki67+ cells in the adult mouse SGZ are thus in the cell cycle or “proliferating” at the time of tissue collection (Mandyam et al., 2007), which is in keeping with the clustering of Ki67+ nuclei seen in the SGZ of both Sham and LFPI mice **(Fig. 1F, 1Gi)**. Stereological assessment of Ki67+ SGZ cells mice 3 dpi revealed an effect of injury **(Fig. 2A)**, with ~45% more Ki67+ SGZ cells in LFPI vs. Sham mice ([statistics for this and all subsequent measures provided in](https://drive.google.com/open?id=1y5qjkG0rUS1P3cbiRrgFePM4lsUDEF2fMch4SbJnkFw) Tables S1 and S2). As the DG, and thus the SGZ, varies along its longitudinal axis in regard to afferents, efferents, and function (Scharfman, 2011; Wu et al., 2015; Levone et al., 2020), Ki67+ cells were also quantified in the anterior vs. posterior SGZ with the division defined as −2.60mm relative to Bregma (Tanti and Belzung, 2013). Similar to the analysis on the entire SGZ **(Fig. 2A)**, analysis of Ki67+ cell number in the anterior and posterior SGZ 3 dpi revealed an effect of injury, with ~40% **(Fig. 2B)** and ~50% more Ki67+ cells **(Fig. 2C)**, respectively, in the ipsilateral neurogenic SGZ of LFPI vs. Sham mice.

**Figure 2.**
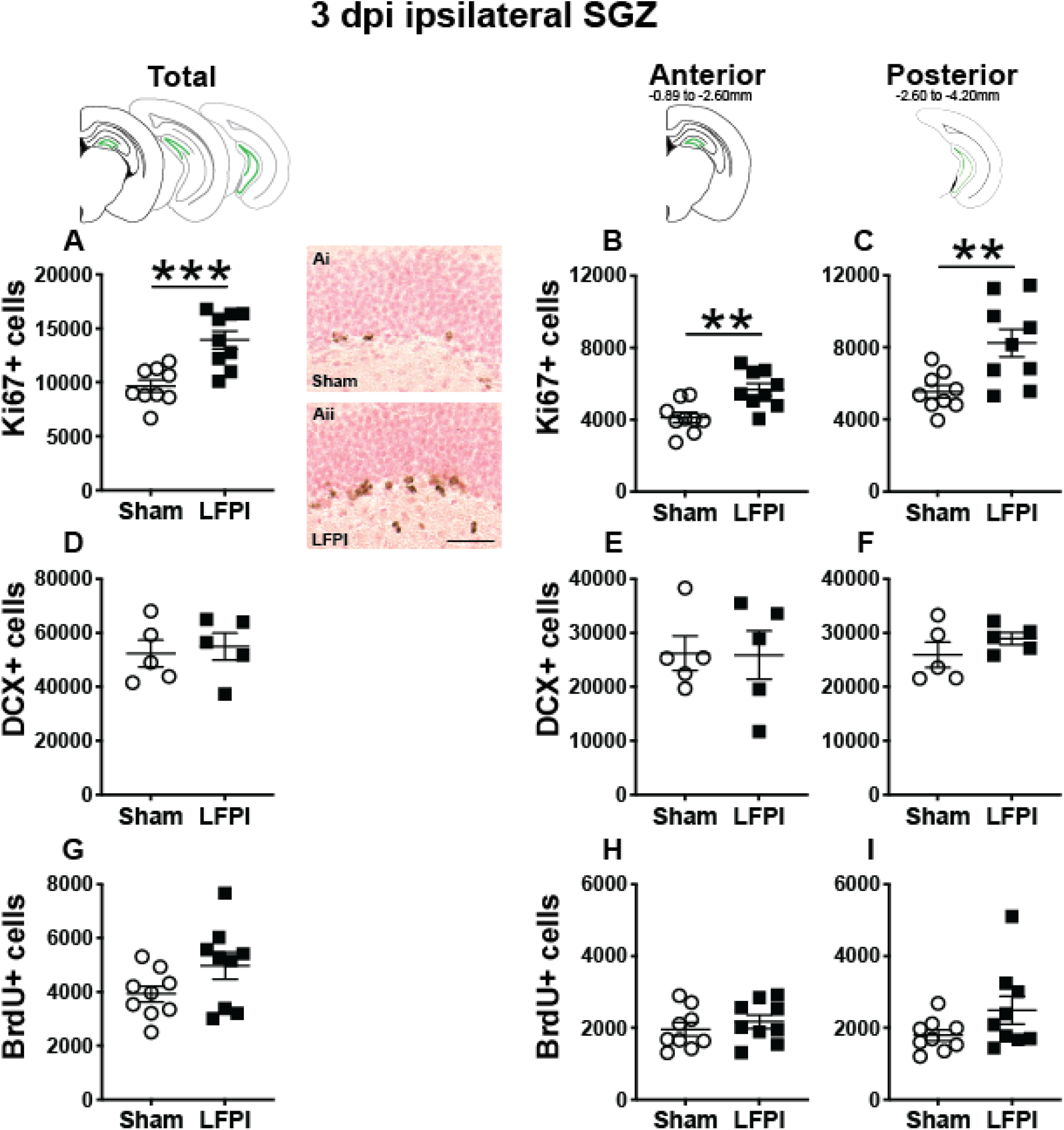
Relative to Sham, LFPI increases the number of Ki67-immunoreactive (Ki67+) proliferating cells in the ipsilateral mouse subgranular zone (SGZ) 3 dpi. As in Fig. 1D, green lines in schematics (top row) indicate these measures were taken in the ipsilateral SGZ/GCL. Stereological quantification of Ki67+ **(A-C**; Sham n=9, LFPI n=9), DCX+ **(D-F**; Sham n=5, LFPI n=5), and BrdU+ **(G-I**; Sham n=9, LFPI n=9)cells in the SGZ (Ki67, BrdU) and SGZ/GCL (DCX). Immunopositive cells were quantified considering immunopositive cells in the SGZ across the entire longitudinal axis (see sample representative schematics above **A, D, G**), and also divided into anterior DG (see sample schematic above **B, E, H**) and posterior DG (see sample schematic above **C, F, I**), operationally-defined as Bregma levels −0.92 to −2.6; and −2.6 to −3.97, respectively. Representative photomicrographs of Sham **(Ai)** and LFPI **(Aii)** Ki67-stained tissue are shown alongside quantification of total Ki67+ cells. Scale bar = 50 μm. One way ANOVA, **p<0.01, ***p<0.001.

#### 3.2.2 Three dpi, the number of DCX+ neuroblasts/immature neurons in the ipsilateral mouse SGZ/GCL is similar between Sham and LFPI mice

The SGZ/GCL of Sham and LFPI mice was examined for cells expressing DCX, a microtubule-associated protein expressed in late progenitor cells (neuroblasts) and immature neurons (Francis et al., 1999; Nacher et al., 2001; Couillard-Despres et al., 2005; La Rosa et al., 2019). DCX+ cells were quantified in both SGZ and GCL as DCX+ cells with neurogenic potential are clearly identifiable in both these DG subregions in control tissue. In the SGZ/GCL of Sham and LFPI mice collected 3 dpi, there was no effect of injury on total DCX+ cell number **(Fig. 2D)**. When the SGZ/GCL in Sham and LFPI mice 3 dpi was divided into anterior and posterior sections, there was also no effect of injury on DCX+ cell number in either the anterior **(Fig. 2E)** or posterior **(Fig. 2F)** bin.

#### 3.2.3 Three dpi, the number of proliferating cells (cells in S phase of the cell cycle) in the ipsilateral mouse SGZ is similar between Sham and LFPI mice

Finally, the SGZ of Sham and LFPI mice was analyzed for cells immunopositive for BrdU, a thymidine analog administered to all mice 3 dpi. BrdU integrates into the DNA of cells in the S phase of the cell cycle at the time of BrdU injection. The short *in vivo* bioavailability of BrdU in mice (<15 min, Mandyam et al., 2007) allows a “pulse” labeling of cycling cells and the ability to track them and their progeny over time. In the ipsilateral SGZ of Sham and LFPI mice collected 3 dpi (2 hrs post-BrdU injection, Short-Term group, **Fig. 1A, 1F**), there was no effect of injury **(Fig. 2G)**. Parcellation of Sham and LFPI BrdU+ SGZ cell counts into anterior and posterior DG bins also showed no effect of injury (**Fig. 2H**, **2I**). While both Ki67 and short-term BrdU were examined as indices of proliferation in the SGZ, they do not represent the same cell population; exogenous BrdU “pulse” labels cells in S-phase, while Ki67 is an endogenous marker of cells in G1, S, G2, and M phase. Thus, despite the lack of change in SGZ BrdU+ cell number between Sham and LFPI mice (which suggests the number of cells in S phase does not change 3 dpi), the greater number of Ki67+ cells in LFPI vs. Sham mice suggests the number of proliferating cells in the entire cell cycle is increased in the ipsilateral SGZ of LFPI mice 3 dpi relative to Sham.

### 3.3 Three dpi: neurogenesis indices in the contralateral DG neurogenic regions

Our main focus for this study was comparing indices of proliferation and neurogenesis in neurogenic regions in the hemisphere ipsilateral to the injury in Sham and LFPI mice, in keeping with the common approach in studies of unilateral injury. However, as the left and right mouse DG are neuroanatomically connected, and as some TBI studies examine or even compare neurogenesis indices between the ipsilateral and contralateral hemispheres (Tran et al., 2006; Gao et al., 2008; Blaiss et al., 2011; Zhou et al., 2012; Hood et al., 2018), we also quantified these indices in the SGZ contralateral to the injury in Sham and LFPI mice.

#### 3.3.1 Three dpi, Ki67+ cell number is similar in the contralateral SGZ between Sham and LFPI mice

In contrast to the Ki67+ cell results from the ipsilateral hemisphere (where there are more Ki67+ cells in LFPI vs. Sham SGZ 3 dpi, **Fig. 2A-C**), in the contralateral SGZ 3 dpi there was no effect of injury on total Ki67+ cell number **(Fig. S1A)** and no effect of injury on Ki67+ cell number in either the anterior or posterior contralateral SGZ **(Fig. S1 B-C)**.

#### 3.3.2 Three dpi, DCX+ cell number is similar in the contralateral SGZ/GCL between Sham and LFPI mice

Similar to the DCX+ cell results from the ipsilateral hemisphere **(Fig. 2D-F)**, in the contralateral SGZ/GCL there was no effect of injury on total DCX+ cell number 3 dpi **(Fig. S1D)** and no effect of injury on DCX+ cell number in either the anterior or posterior contralateral SGZ/GCL **(Fig. S1 E-F)**.

#### 3.3.3 Three dpi, BrdU+ cell number (cells in S phase of the cell cycle) is similar in the contralateral SGZ between Sham and LFPI mice

Similar to the BrdU+ cell results from the ipsilateral hemisphere **(Fig. 2G-I)**, in the contralateral SGZ there was no effect of injury on total BrdU+ cell number 3 dpi **(Fig. S1G)** and no effect of injury on BrdU+ cell number in either the anterior or posterior contralateral SGZ **(Fig. S1 H-I)**.

### 3.4 Three dpi: proliferation indices in the ipsilateral and contralateral DG non-neurogenic regions

In naive rodents, new neurons primarily emerge from the main DG neurogenic region: the SGZ/inner portion of the GCL; it is rare for other DG subregions (hilus, oGCL, molecular layer) to give rise to new neurons. However, in naive rodents, these “non-neurogenic” subregions contain proliferating cells (García-Martinez et al., 2020). Notably, proliferation in these non-neurogenic DG regions is often different in injured vs. sham rodents in both the ipsilateral and contralateral hemispheres (Cho et al., 2015; Du et al., 2017; Neuberger et al., 2017). While these injury-induced changes in proliferation in non-neurogenic DG regions may reflect division of precursors of glia, not neurons (Avendaño and Cowan, 1979; Dragunow et al., 1990; Hailer et al., 1999; Blümcke et al., 2001; Littlejohn et al., 2020), the impact of glia on DG structure and function stress the importance of understanding how mTBI influences these measures. Therefore, 3 dpi the ipsilateral and contralateral hilus, oGCL, and molecular layer from Sham and LFPI mice were assessed for indices of proliferation (Ki67+ and BrdU+ cell number). For DCX+ neuroblasts/immature neurons, the oGCL DCX+ cells were considered as part of the SGZ/GCL analysis (**Figs. 2D-F**, **S1D-F**; see sections 3.2.2 and 3.3.2); due to the presence of faint or ambiguous DCX+ cells in the hilus and molecular layer with this and other DCX+ antibodies in both experimental and control groups, DCX+ cells were not quantified in the hilus or molecular layer.

#### 3.4.1 Three dpi, there are more proliferating (Ki67+ and BrdU+ cells) in ipsilateral non-SGZ DG subregions in LFPI mice relative to Sham mice

In the ipsilateral hilus 3 dpi, there was an effect of injury on total Ki67+ cells **(Fig. 3A)** and BrdU+ cells **(Fig. 3D)**, with 135% and 130% more Ki67+ and BrdU+ cells, respectively, in LFPI vs. Sham mice. In the ipsilateral hemisphere, parcellation of Sham and LFPI BrdU+ hilus cell counts into anterior and posterior DG bins revealed significant effects of injury for both Ki67+ and BrdU+ cells in the anterior and posterior hilus. In the ipsilateral hemisphere, LFPI mice had 100% more anterior Ki67+ cells **(Fig. 3B)**, 150% more posterior Ki67+ cells **(Fig. 3C)**, 90% more anterior BrdU+ cells **(Fig. 3E)**, and 150% more posterior BrdU+ cells **(Fig. 3F)** in the hilus vs. Sham mice. In the ipsilateral oGCL 3 dpi, there was also an effect of injury on Ki67+ cells, with 275% more total **(Fig. 3G)**, 260% more anterior **(Fig. 3H)**, and ~290% more posterior **(Fig. 3I)** Ki67+ cells in LFPI vs. Sham mice. However, in the ipsilateral oGCL 3 dpi, there was no effect of injury on total BrdU+ oGCL cells **(Fig. 3J)** or on anterior or posterior BrdU+ oGCL cells **(Fig. 3 K-L)**. In the ipsilateral molecular layer 3 dpi, there was an effect of injury on total Ki67+ **(Fig. 3M)** and BrdU+ (**Fig. 3P**) cells, with 240% and 160% more Ki67+ and BrdU+ cells, respectively, in LFPI vs. Sham mice. In the ipsilateral parcellation of Sham and LFPI BrdU+ molecular layer cell counts into anterior and posterior DG bins revealed significant effects of injury with 250% more anterior Ki67+ **(Fig. 3N)**, 230% more posterior Ki67+ **(Fig. 3O)**, 80% more anterior BrdU+ **(Fig. 3Q)**, and 270% more posterior BrdU+ cells **(Fig. 3R)** cells in the molecular layer of LFPI vs. Sham mice. These results show 3 dpi there is increased proliferation in the ipsilateral dentate gyrus in DG subregions that are classically believed to be non-neurogenic.

**Figure 3.**
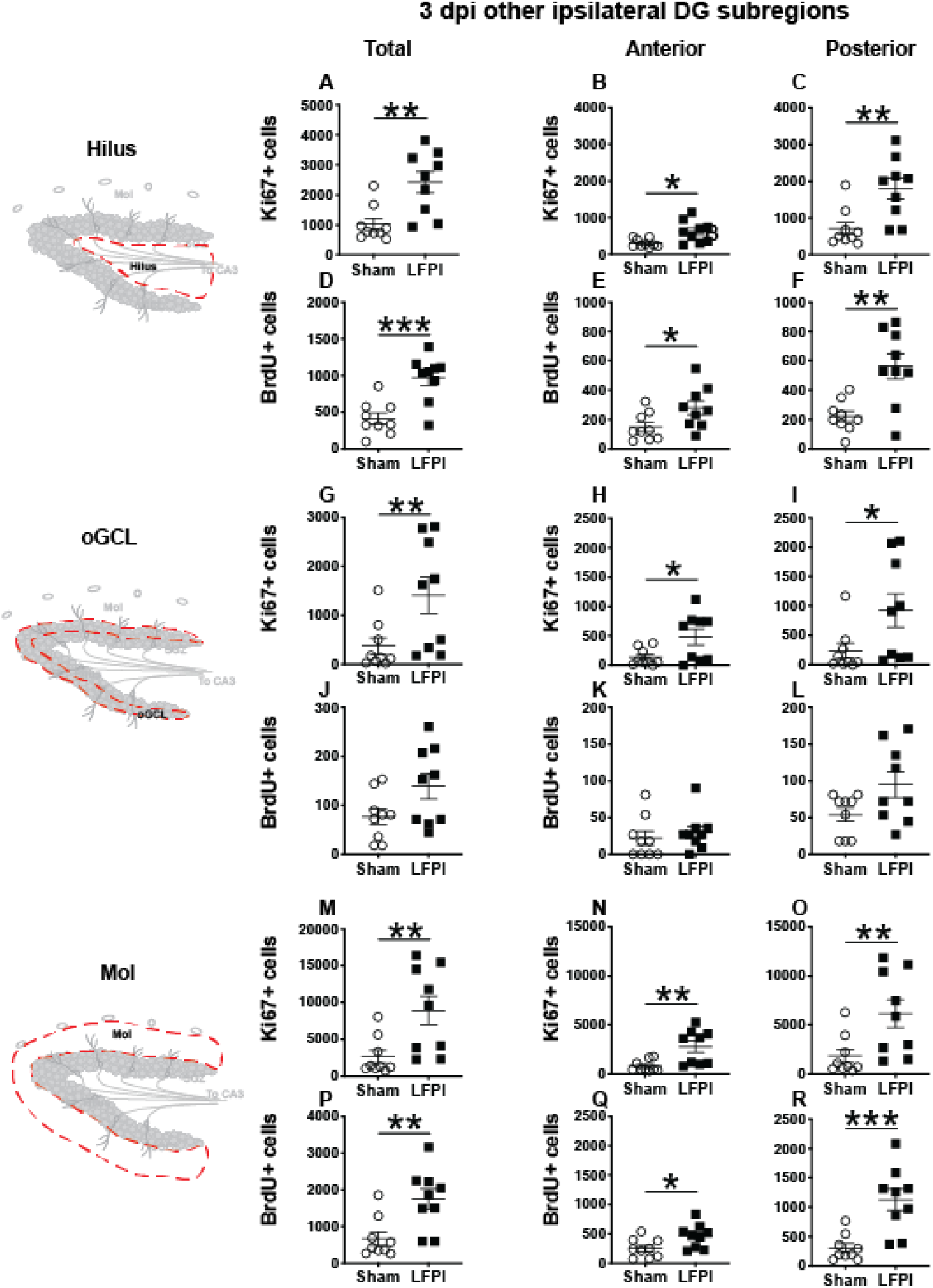
Relative to Sham, LFPI increases the number of Ki67+ and BrdU+ proliferating cells in the ipsilateral DG hilus and molecular layer (Mol) 3 dpi. Stereological quantification of Ki67+ **(A-C, G-I, M-O**; Sham n=9, LFPI n=9)and BrdU+ **(D-F, J-L, P-R**; Sham n=9, LFPI n=9)cells in the hilus (**A-F**; red dotted line region, top-left schematic), outer granule cell layer (**G-L**; red dotted line region, middle-left schematic), and molecular layer (**M-R**; red dotted line region, bottom-left schematic). Immunopositive cells were quantified across the entire longitudinal axis **(A, D, G, J, M, P)**, and also broken up into anterior **(B, E, H, K, N, Q)** and posterior **(C, F, I, L, O, R)** bins, operationally defined as Bregma levels −0.92 to −2.6; and −2.6 to −3.97, respectively. One way ANOVA, *p<0.05, **p<0.01, ***p<0.001.

#### 3.4.2 Three dpi, there are more proliferating Ki67+ cells in some contralateral non-SGZ DG subregions in LFPI mice relative to Sham mice

In the contralateral hilus 3 dpi, there was an effect of injury on Ki67+ cells, with 65% more total **(Fig. S2A)** and 50% more posterior **(Fig. S2C)** Ki67+ cells in LFPI vs. Sham mice; the number of Ki67+ cells in the contralateral anterior hilus was similar between Sham and LFPI mice **(Fig. S2B)**. There was no effect of injury on total **(Fig. S2D)** or anterior or posterior **(Fig. S2 E-F)** BrdU+ cell number in the contralateral hilus 3 dpi. In the contralateral oGCL 3 dpi, there was an effect of injury on Ki67+ cells, with 475% more total **(Fig. S2G)** and 520% more posterior **(Fig. S2I)** Ki67+ cells in LFPI vs. Sham mice; as in the contralateral hilus, Ki67+ cell number in the contralateral anterior oGCL was similar between Sham and LFPI mice 3 dpi **(Fig. S2H)**. There was no effect of injury on total **(Fig. S2J)** or anterior **(Fig. S2K)** BrdU+ cells, but there were 75% fewer BrdU+ cells in the contralateral posterior oGCL **(Fig. S2L)** in LFPI vs. Sham mice 3 dpi. In the contralateral molecular layer 3 dpi, there was no effect of injury on Ki67+ cell number (total **Fig. S2M**; anterior and posterior **Fig. S2 N-O**) or BrdU+ cell number (total **Fig. S2P**, anterior and posterior **Fig. S2 Q-R)** 3 dpi. These results show 3 dpi there is increased proliferation in the contralateral hilus and oGCL in LFPI vs. Sham mice, suggesting the influence of mTBI on proliferation is not restricted to the ipsilateral neurogenic and non-neurogenic regions. As in the ipsilateral 3 dpi analyses, in the contralateral 3 dpi analysis there is a disconnect between the impact of mTBI on the number of cells in S phase (BrdU+ cells, which are similar between Sham and LFPI in all contralateral DG subregions, with the exception of fewer BrdU+ cells in the oGCL in LFPI vs. Sham mice) and on cells in the entire cell cycle (Ki67+ cells, which are similar between Sham and LFPI in the contralateral SGZ but increased in the contralateral hilus and oGCL in LFPI vs. Sham mice).

### 3.5 Seven dpi: neurogenesis indices in the ipsilateral DG neurogenic regions

The same markers examined in ipsilateral 3 dpi DG were also analyzed in the ipsilateral DG of brains collected 7 dpi **(Fig. 4)**. Since all mice received a BrdU injection 3 dpi, by 7 dpi these BrdU+ cells represent cells that were in S-phase of the cell cycle 4 days prior, but may no longer be actively proliferating (Cameron and McKay, 2001; Dayer et al., 2003; Mandyam et al., 2007).

**Figure 4.**
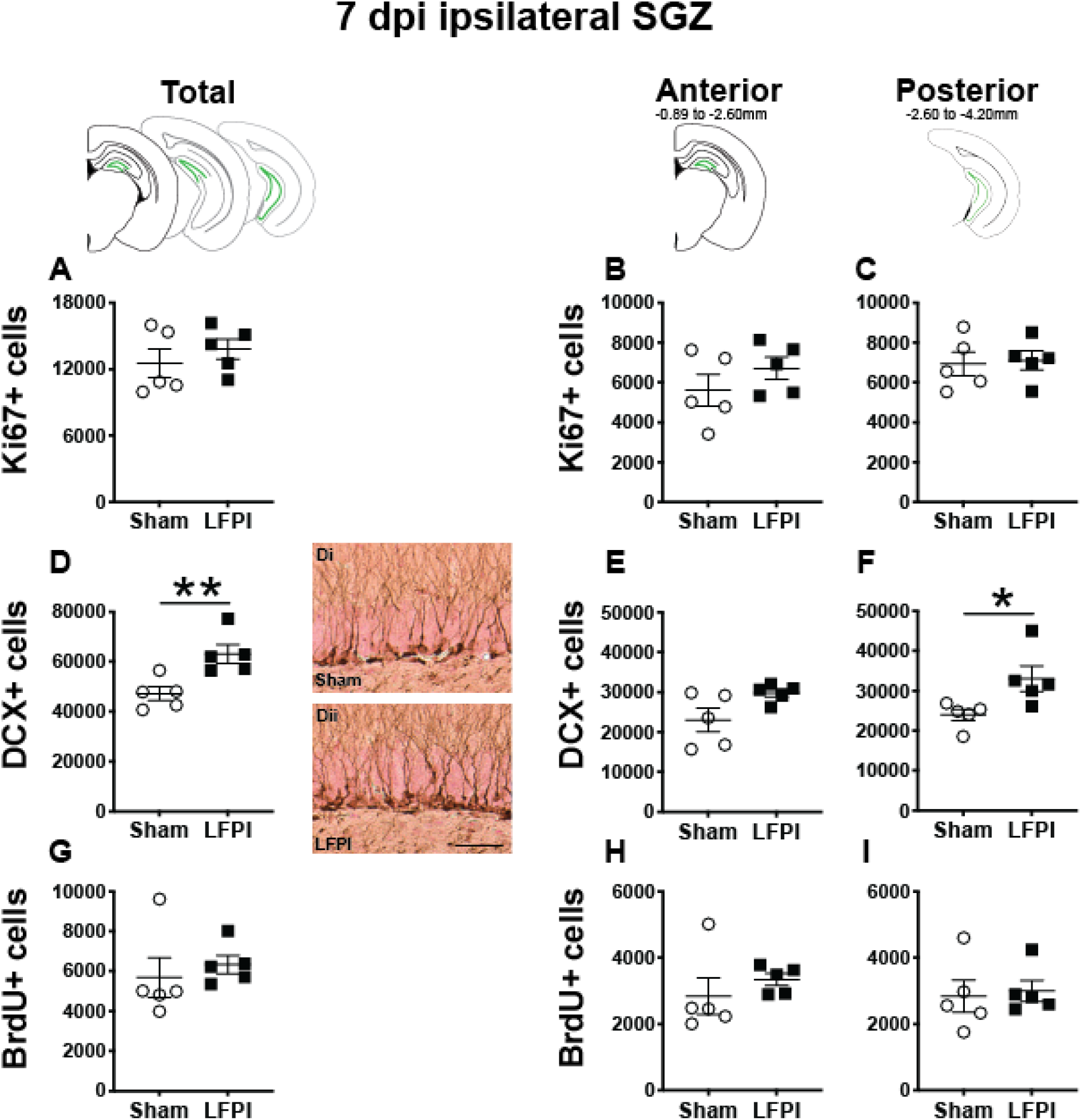
Relative to Sham, LFPI increases the number of DCX+ neuroblasts/immatures neurons in the ipsilateral mouse SGZ/GCL 7 dpi. Green lines in schematics (top row) indicate these measures were taken in the ipsilateral SGZ/GCL. Stereological quantification of Ki67+ **(A-C**; Sham n=5, LFPI n=5), DCX+ **(D-F**; Sham n=5, LFPI n=5), and BrdU+ **(G-I**; Sham n=5, LFPI n=5)cells in the SGZ (Ki67, BrdU) and GCL (DCX). Immunopositive cells were quantified across the entire longitudinal axis **(A, D, G)**, and also broken up into anterior **(B, E, H)** and posterior **(C, F, I)** bins, operationally defined as Bregma levels −0.92 to −2.6; and −2.6 to −3.97, respectively. Representative photomicrographs of Sham **(Di)** and LFPI **(Dii)** DCX-stained tissue are shown alongside quantification of total DCX+ cells. Scale bar = 50 μm. One way ANOVA, **p<0.01.

#### 3.5.1 Seven dpi, there are the same number of Ki67+ proliferating cells in the ipsilateral SGZ in LFPI and Sham mice

In the ipsilateral SGZ of Sham and LFPI mice 7 dpi, there was no effect of injury on total Ki67+ cell number **(Fig. 4A)**. When divided into anterior/posterior bins, there was no effect of injury on Ki67+ cell number in either the anterior **(Fig. 4B)** or posterior **(Fig 4C)** ipsilateral SGZ 7 dpi.

#### 3.5.2 Seven dpi, there are more DCX+ neuroblasts/immature neurons in the ipsilateral mouse SGZ/GCL in LFPI mice relative to Sham mice

In the ipsilateral SGZ/GCL of Sham and LFPI mice 7 dpi, there was an effect of injury **(Fig. 4D)**, with ~35% more total DCX+ SGZ/GCL cells in LFPI vs. Sham mice. When divided into anterior/posterior bins, there was no effect of injury on DCX+ cell number in the ipsilateral anterior SGZ/GCL **(Fig 4E)**, but there was an effect of injury on DCX+ cell number in the ipsilateral posterior SGZ/GCL **(Fig. 4F)** with ~40% more DCX+ SGZ/GCL cells in LFPI vs. Sham mice 7 dpi.

#### 3.5.3 Seven dpi, the number of BrdU+ cells (cells in S phase 4 days prior) in the ipsilateral mouse SGZ is similar between Sham and LFPI mice

In the ipsilateral SGZ of Sham and LFPI mice 7 dpi, there was no effect of injury on total BrdU+ cell number **(Fig. 4G)**. When divided into anterior/posterior bins, there was no effect of injury on BrdU+ cell number in either the anterior **(Fig. 4H)** or posterior **(Fig. 4I)** ipsilateral SGZ 7 dpi.

### 3.6 Seven dpi: neurogenesis indices in the contralateral DG neurogenic regions

#### 3.6.1 Seven dpi, the number of Ki67+ proliferating cells in the contralateral SGZ is similar between Sham and LFPI mice

In the contralateral SGZ of Sham and LFPI mice 7 dpi, there was no effect of injury on total Ki67+ cell number **(Fig. S3A)** or on Ki67+ cell number in the anterior **(Fig. S3B)** or posterior **(Fig. S3C)** contralateral SGZ.

#### 3.6.2 Seven dpi, the number of BrdU+ cells (cells in S phase 4 days prior) in the contralateral SGZ in is similar between Sham and LFPI mice

In the contralateral SGZ of Sham and LFPI mice 7 dpi, there was no effect of injury on total BrdU+ cell number **(Fig. S3A)** or on BrdU+ cell number in either the anterior **(Fig. S3B)** or posterior **(Fig. S3C)** contralateral SGZ.

### 3.7 Seven dpi: proliferation indices in the ipsilateral and contralateral DG non-SGZ regions

#### 3.7.1 Seven dpi, the numbers of Ki67+ proliferating cells and BrdU+ cells (cells in S phase 4 days prior) in the ipsilateral non-SGZ DG subregions are similar in Sham and LFPI mice

In the ipsilateral DG 7 dpi, there was no effect of injury on Ki67+ cell number in the total hilus, oGCL, or Mol **(Fig. 5A, G, M)** or in the anterior or posterior hilus, oGCL, or Mol **(Fig. 5B-C, H-I, N-O)**. Similarly, in the ipsilateral DG 7 dpi, there was no effect of injury on BrdU+ cell number in the total hilus, oGCL, or Mol **(Fig. 5D, J, P)** or in the anterior or posterior hilus, oGCL, or Mol **(Fig. 5E-F, K-L, Q-R)**.

**Figure 5.**
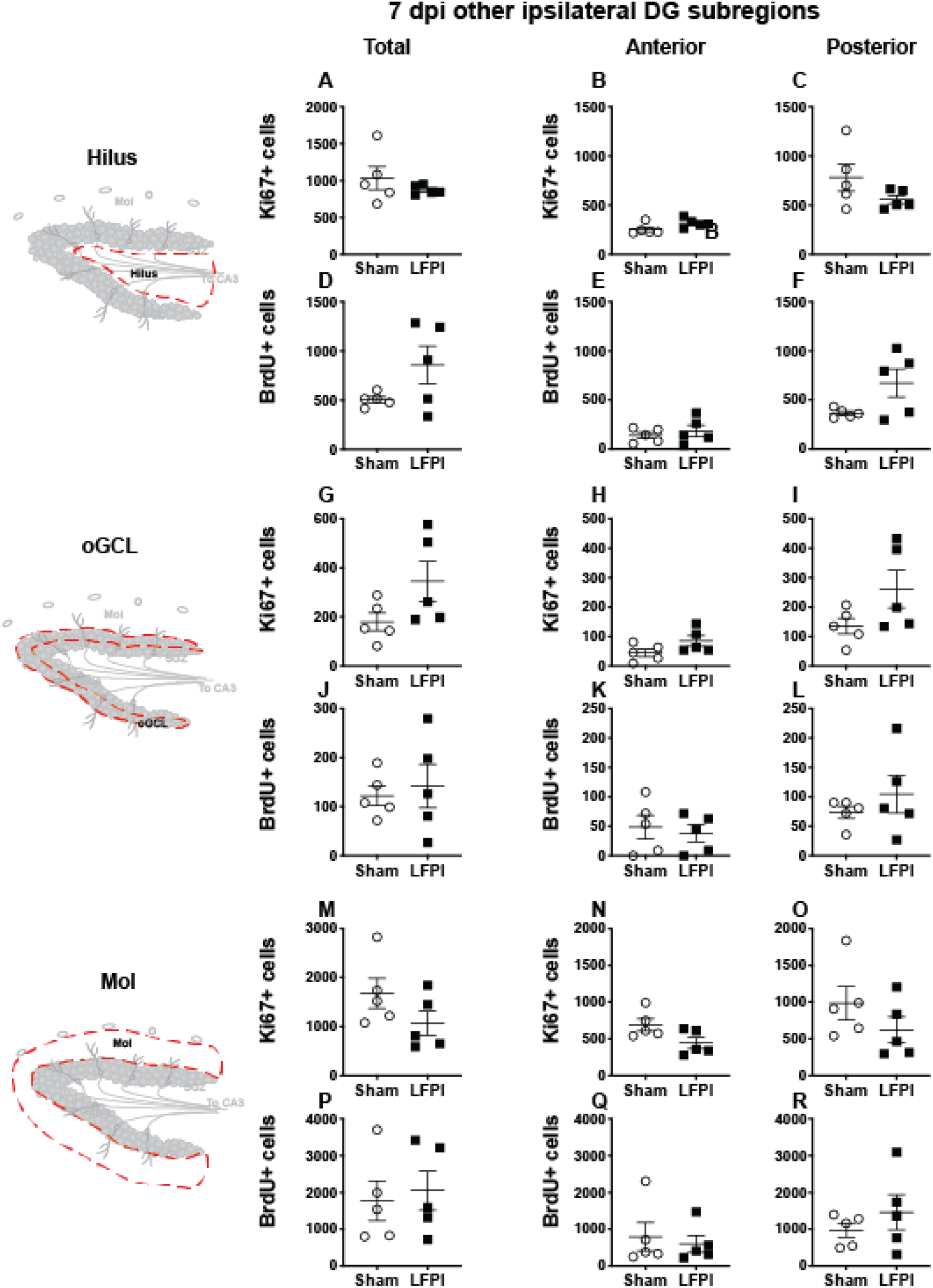
Relative to Sham, LFPI does not change the number of Ki67+ or BrdU+ cells in the ipsilateral DG hilus, oGCL, and Mol 7 dpi. Stereological quantification of Ki67+ **(A-C, G-I, M-O**; Sham n=5, LFPI n=5) and BrdU+ **(D-F, J-L, P-R**; Sham n=5, LFPI n=5)cells in the hilus (**A-F**; red dotted line region, top-left schematic), outer granule cell layer (**G-L**; red dotted line region, middle-left schematic), and molecular layer (**M-R**; red dotted line region, bottom-left schematic). Immunopositive cells were quantified across the entire longitudinal axis **(A, D, G, J, M, P)**, and also broken up into anterior **(B, E, H, K, N, Q)** and posterior **(C, F, I, L, O, R)** bins, operationally defined as Bregma levels −0.92 to −2.6; and −2.6 to −3.97, respectively.

#### 3.7.2 Seven dpi, the numbers of Ki67+ proliferating cells and BrdU+ cells (cells in S phase 4 days prior) in the contralateral non-SGZ DG subregions are similar in Sham and LFPI mice

In the contralateral DG 7 dpi, there was no effect of injury on Ki67+ cell number in the total hilus, oGCL, or Mol **(Fig. S4A, G, M)** or in the anterior or posterior hilus, oGCL, or Mol **(Fig. S4B-C, H-I, N-O)**. Similarly, in the contralateral DG 7 dpi, there was no effect of injury on BrdU+ cell number in the total hilus, oGCL, or Mol **(Fig. S4D, J, P)** or in the anterior or posterior hilus, oGCL, or Mol **(Fig. S4E-F, K-L, Q-R)**.

### 3.8 Thirty-one dpi: neurogenesis indices in the ipsilateral DG neurogenic regions

The same markers examined in 3 and 7 dpi ipsilateral tissue were also analyzed in tissue harvested 31 dpi **(Fig. 4)**. BrdU+ cells 31 dpi reflect cells that were in S-phase of the cell cycle 28 days prior.

#### 3.8.1 Thirty-one dpi, the number of Ki67+ proliferating cells in the ipsilateral mouse SGZ is similar between Sham and LFPI mice

In the ipsilateral SGZ 31 dpi, there was no effect of injury on Ki67+ cell number in the total **(Fig. 6A)** or the anterior or posterior SGZ **(Fig. 6B, C)**. Given the cell cycle length of mouse SGZ cells and the dilution of BrdU that occurs with additional cell division (Cameron and McKay, 2001; Dayer et al., 2003; Mandyam et al., 2007), detectable BrdU+ SGZ cells 31 dpi are unlikely to be still proliferating, and thus here are interpreted as surviving cells.

**Figure 6.**
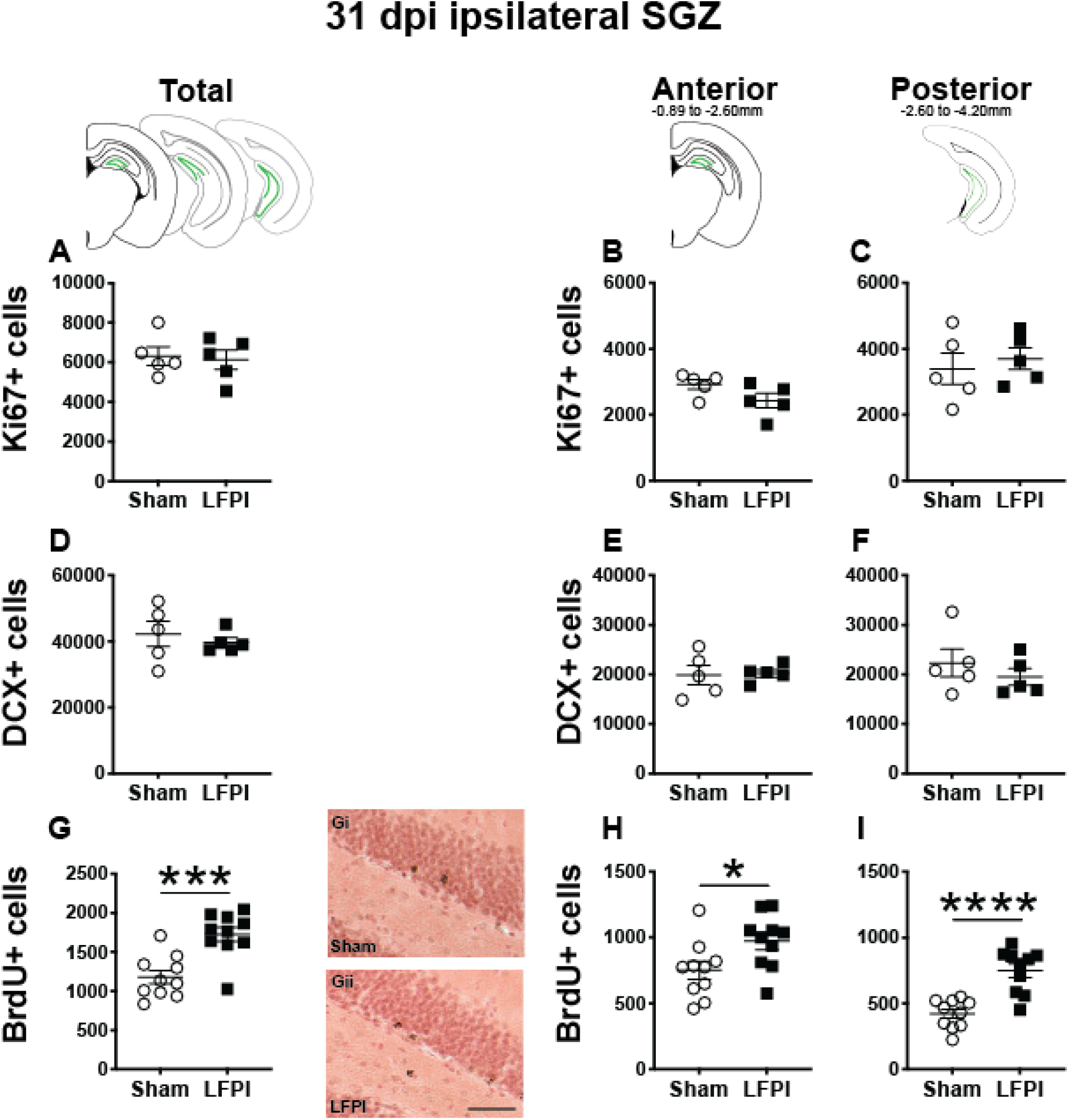
Relative to Sham, LFPI increases the number of BrdU+ surviving cells in the ipsilateral mouse SGZ 31 dpi. Green lines in schematics (top row) indicate these measures were taken in the ipsilateral SGZ/GCL. Stereological quantification of Ki67+ **(A-C**; Sham n=5, LFPI n=5), DCX+ **(D-F**; Sham n=5, LFPI n=5), and BrdU+ **(G-I**; Sham n=10, LFPI n=10)cells in the SGZ (Ki67, BrdU) and GCL (DCX). Immunopositive cells were quantified across the entire longitudinal axis **(A, D, G)**, and also broken up into anterior **(B, E, H)** and posterior **(C, F, I)** bins, operationally defined as Bregma levels −0.92 to −2.6; and −2.6 to −3.97, respectively. Representative photomicrographs of Sham **(Gi)** and LFPI **(Gii)** BrdU-stained tissue are shown alongside quantification of total BrdU+ cells. Scale bar = 50 μm. One way ANOVA, *p<0.05, ***p<0.001, ****p<0.0001.

#### 3.8.2 Thirty-one dpi, the number of DCX+ neuroblasts/immature neurons in the ipsilateral mouse SGZ/GCL is similar between Sham and LFPI mice

In the ipsilateral DG 31 dpi, there was no effect of injury on DCX+ cell number in the total **(Fig. 6A)** or the anterior or posterior SGZ/GCL **(Fig. 6B, C)**.

#### 3.8.3 Thirty-one dpi (28 d post-BrdU injection), the number of BrdU+ cells in the ipsilateral mouse SGZ is greater in LFPI vs. Sham mice

In the ipsilateral DG 31 dpi, there was an effect of injury on BrdU+ cell number in the total SGZ **(Fig. 6G)**, with ~45% more cells in LFPI vs. Sham mice. When divided into anterior/posterior bins, there was also an effect of injury with ~30% **(Fig. 6H)** and ~75% more BrdU+ cells **(Fig. 6I)**, respectively, in LFPI vs. Sham mice **(Fig. 6B, C)**. These data suggest that all the number of cells in S-phase is unchanged when BrdU is first injected **(Fig. 2G-I)**, survival of those BrdU+ cells and/or their progeny is increased in LFPI mice by 31 dpi vs. Sham mice.

### 3.9 Thirty-one dpi: neurogenesis indices in the contralateral DG neurogenic regions

#### 3.9.1 Thirty-one dpi, the number of Ki67+ proliferating cells in the contralateral SGZ is similar between Sham and LFPI mice

In the contralateral SGZ of Sham and LFPI mice 31 dpi, there was no effect of injury on total Ki67+ cell number **(Fig. S5A)** or on Ki67+ cell number in the anterior **(Fig. S5B)** or posterior **(Fig. S5C)** contralateral SGZ.

#### 3.9.2 Thirty-one dpi, the number of DCX+ immature neurons/neuroblasts in the contralateral SGZ/GCL is similar between Sham and LFPI mice

In the contralateral SGZ/GCL of Sham and LFPI mice 31 dpi, there was no effect of injury on total DCX+ cell number **(Fig. S5D)** or on DCX+ cell number in the anterior **(Fig. S5E)** or posterior **(Fig. S5F)** contralateral SGZ.

#### 3.9.3 Thirty-one dpi (28 d post-BrdU injection), the number of BrdU+ cells in the contralateral SGZ is similar between Sham and LFPI mice

In the contralateral SGZ of Sham and LFPI mice 31 dpi, there was no effect of injury on total BrdU+ cell number **(Fig. S5G)** or on BrdU+ cell number in the anterior **(Fig. S5H)** or posterior **(Fig. S5I)** contralateral SGZ.

### 3.10 Thirty-one dpi: proliferation and cell survival indices in the ipsilateral and contralateral D G non-neurogenic regions

#### 3.10.1 Thirty-one dpi, the number of Ki67+ proliferating cells in the ipsilateral non-SGZ DG subregions are similar in Sham and LFPI mice, but the numbers of surviving BrdU+ cells in the ipsilateral non-SGZ DG subregions are higher in LFPI vs. Sham mice

In the ipsilateral DG 31 dpi, there was no effect of injury on Ki67+ cell number in the total hilus, oGCL, or Mol (Fig. 7A, G, M)or in the anterior or posterior hilus, oGCL, or Mol (Fig. 7B-C, H-I, N-O). However similar to the ipsilateral SGZ 31 dpi, in the ipsilateral DG 31 dpi, there was an effect of injury on BrdU+ cell number in the total hilus, oGCL, and Mol **(Fig. 7D, J, P)** with ~460%, 75%, and 230% more BrdU+ cells, respectively, in LFPI vs Sham mice. When divided into anterior/posterior bins, there was an effect of injury in the anterior and posterior hilus **(Fig. 7E-F)**, posterior oGCL **(Fig. 7L)**, and anterior and posterior molecular layer **(Fig. 7Q-R)** with 460%, 470%, 85%, 210%, and 260% more BrdU+ cells in these regions in LFPI vs. Sham mice 31 dpi. Together, these data suggest that more BrdU+ cells in other non-SGZ subregions survive out to 31 dpi in LFPI vs. Sham mice.

**Figure 7.**
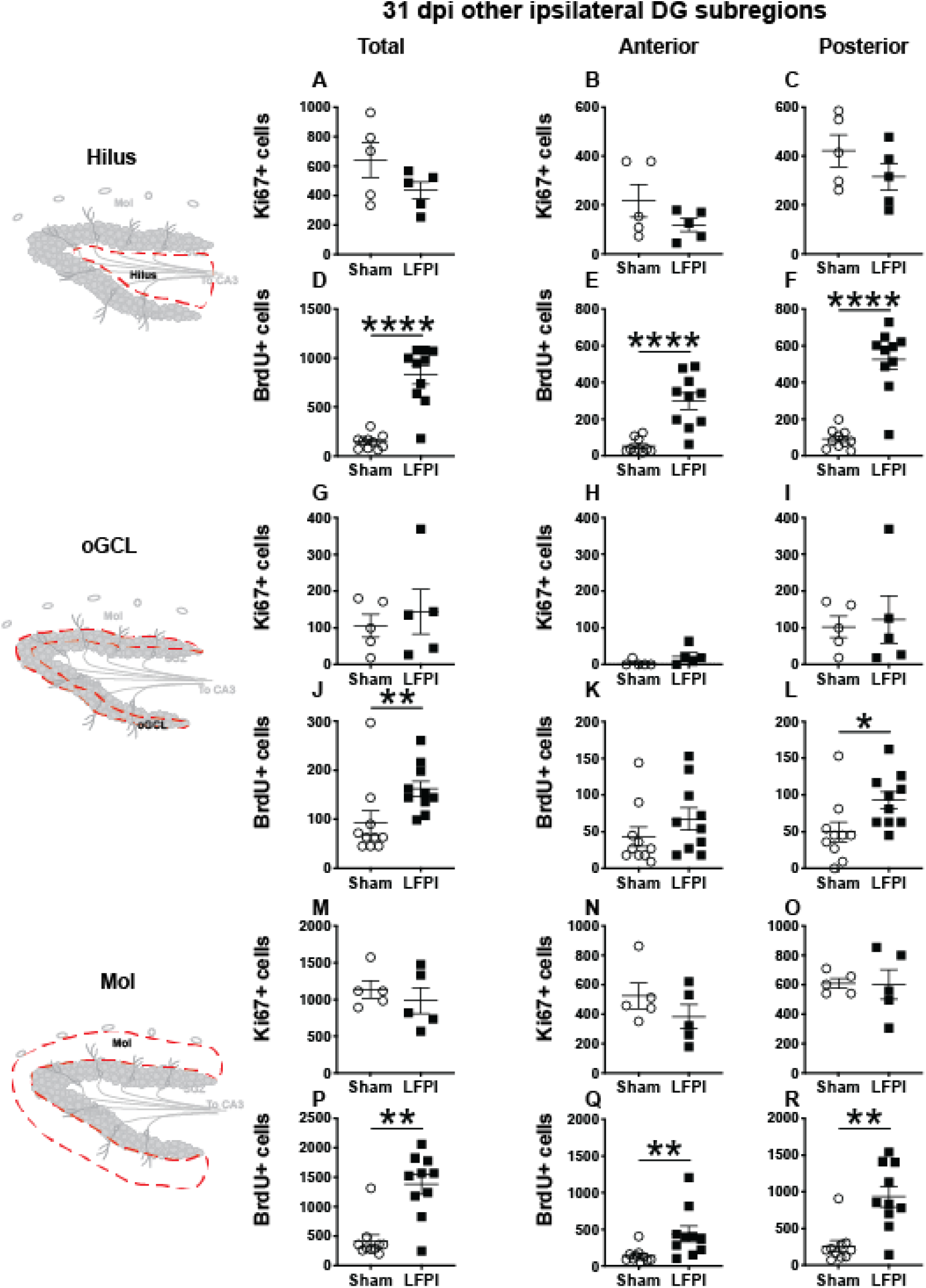
Relative to Sham, LFPI increases the number of BrdU+ surviving cells in the ipsilateral DG hilus, posterior oGCL, and Mol 31 dpi. Stereological quantification of Ki67+ **(A-C, G-I, M-O**; Sham n=5, LFPI n=5)and BrdU+ **(D-F, J-L, P-R**; Sham n=10, LFPI n=10)cells in the hilus (**A-F**; red dotted line region, top-left schematic), outer granule cell layer (**G-L**; red dotted line region, middle-left schematic), and molecular layer (**M-R**; red dotted line region, bottom-left schematic). Immunopositive cells were quantified across the entire longitudinal axis **(A, D, G, J, M, P)**, and also broken up into anterior **(B, E, H, K, N, Q)** and posterior **(C, F, I, L, O, R)** bins, operationally defined as Bregma levels −0.92 to −2.6; and −2.6 to −3.97, respectively. One way ANOVA, *p<0.05, **p<0.01, ****p<0.0001.

#### 3.10.2 Thirty-one dpi, the numbers of Ki67+ proliferating cells and BrdU+ cells in the contralateral non-SGZ DG subregions are similar in Sham and LFPI mice

In the contralateral DG 31 dpi, there was no effect of injury on Ki67+ cell number in the total hilus, oGCL, or Mol **(Fig. S6A, G, M)** or in the anterior or posterior hilus, oGCL, or Mol **(Fig. S6B-C, H-I, N-O)**. Similarly, and in contrast to the greater number of surviving BrdU+ cells in the ipsilateral hemisphere in LFPI vs. Sham mice, in the contralateral DG 31 dpi, there was no effect of injury on BrdU+ cell number in the total hilus, oGCL, or Mol **(Fig. S6D, J, P)** or in the anterior or posterior hilus, oGCL, or Mol **(Fig. S6E-F, K-L, Q-R)**.

### 3.11 DG GCL volume 3, 7, and 31 days after Sham or LFPI

Given that in the ipsilateral hemisphere LFPI mice had greater proliferation 3 dpi, more immature/neuroblasts 7 dpi, and more surviving BrdU+ cells 31 dpi relative to Sham mice **(Fig. 8)**, it is important to see if these cell number changes influence total volume of the GCL. In fact, assessing GCL volume is critical for interpretation of cell number changes (McDonald and Wojtowicz, 2005; Harburg et al., 2007; Brummelte and Galea, 2010). The results of the Cavalieri Estimator to measure ipsilateral and contralateral GCL volume 3, 7, and 31 dpi are provided in **Table 1**. In brief, there was no effect of injury on GCL volume at any time point in either the ipsilateral or contralateral hemisphere. These volume data suggest that the change in neurogenesis is a true change in the underlying process, rather than a matter of scale to match volume changes.

**Table 1.** Granule cell layer (GCL) volume in the ipsilateral and contralateral hemispheres, 3, 7, and 31 d ays post-injury (dpi). Mean ipsilateral and contralateral GCL volume values in Sham and LFPI mice at 3 different time points, shown in mm^3^.

| Time point | Ipsilateral GCL volume (mm<sup>3</sup>) |  |  |  | Contralateral GCL volume (mm<sup>3</sup>) |  |  |  |
| --- | --- | --- | --- | --- | --- | --- | --- | --- |
|  | Sham |  | LFPI |  | Sham |  | LFPI |  |
|  | <u>Mean</u> | <u>SEM</u> | <u>Mean</u> | <u>SEM</u> | <u>Mean</u> | <u>SEM</u> | <u>Mean</u> | <u>SEM</u> |
| 3 dpi | 0.627 | 0.016 | 0.582 | 0.016 | 0.632 | 0.021 | 0.605 | 0.019 |
| 7 dpi | 0.658 | 0.039 | 0.623 | 0.033 | 0.678 | 0.015 | 0.659 | 0.015 |
| 31 dpi | 0.602 | 0.020 | 0.543 | 0.030 | 0.591 | 0.025 | 0.548 | 0.017 |

**Figure 8.**
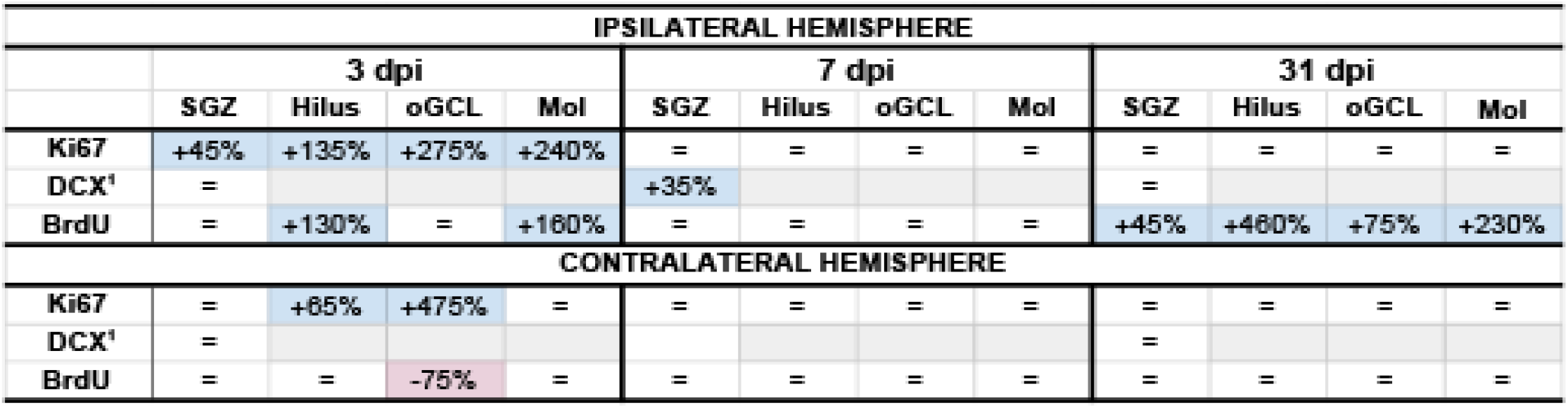
Summary of neurogenesis changes in LFPI vs. Sham mice at three time points post-injury. Percent change in LFPI vs. Sham number of cells immunoreactive for Ki67, DCX, and BrdU in the ipsilateral (top) and contralateral (bottom) dentate gyrus (DG) subregions 3 dpi (left column), 7 dpi (middle column), and 31 dpi (right column). These summary percentages reflect data presented in Figs. 2-7 and Figs. S1-S6. =: not changed vs. Sham. Light-blue shading: increased in LFPI vs. Sham. Rose shading: decreased in LFPI vs. Sham. White: not determined. Light-gray shading: not part of this study. ^1^: quantified in both SGZ and GCL. Mol: molecular layer. ND, not determined. oGCL: outer granule cell layer. SGZ: subgranular zone.

## DISCUSSION

In the preclinical TBI literature - even within the minority of studies focused specifically on mild injuries - there is disagreement about if and how mTBI changes the process of DG neurogenesis. Here we utilized a clinically-relevant rodent model of mTBI (LFPI), gold-standard neurogenesis markers and quantification approaches, three time points post-injury, and analysis of defined DG subregions to generate a comprehensive picture of how mTBI affects the process of adult hippocampal DG neurogenesis. We report five major findings, which are also summarized in **Figure 8**. In the ipsilateral hemisphere classical neurogenic DG regions (SGZ/GCL), LFPI leads to 1) a transient increase in proliferating cells 3 dpi; 2) a transient increase in neuroblasts/immature neurons 7 dpi; and 3) sustained survival of BrdU+ cells 31 dpi, all relative to Sham. In ipsilateral hemisphere DG subregions typically considered to be non-neurogenic (hilus, oGCL, molecular layer), LFPI leads to 4) a transient increase in proliferating cells 3 dpi and 5) sustained survival of BrdU+ cells 31 dpi, both relative to Sham. Our ipsilateral data are in line with the large fraction of literature that reports increased DG neurogenesis in other and more severe models of TBI. However, our data in this mTBI model importantly add temporal, stage-specific, and DG subregional resolution to the literature. Below we discuss these findings in the context of the current literature, consider several interpretations of these data, and mention what they mean for efforts to harness the regenerative capacity of adult-generated DG neurons as a treatment for mTBI-induced cognitive changes.

To gain temporal resolution on how mTBI influences the process of DG neurogenesis, we first examined tissue 3 dpi to a) avoid the immediate mechanical effects of the LFPI, b) give the brain time to mount a response to the injury, and c) select a time point frequently represented in the TBI literature. We found more Ki67+ cells in the ipsilateral SGZ 3 dpi, consistent with the idea that there is increased proliferation in the ipsilateral DG neurogenic niche after TBI (Dash et al., 2001; Chirumamilla et al., 2002; Villasana et al., 2015, 2019). Brain injury-induced increased DG proliferation at this short time point may be due to a variety of factors, from proliferation of microglia or astrocytes subsequent to inflammation (Rola et al., 2006; Piao et al., 2013; Chiu et al., 2016; McAteer et al., 2016; Bedi et al., 2018; Krukowski et al., 2018; Wang et al., 2019) or proliferation of neural stem cells and progenitors due to abnormal network activity. Careful readers may have noticed more Ki67+ cells **(Fig. 2A-C)** but no change in BrdU+ cell number **(Fig. 2 G-I)** in LFPI vs. Sham mice 3 dpi **(Fig. 8)**. This apparent discrepancy between these markers of proliferating cells can be accounted for by their distinct uses: exogenous BrdU is incorporated only by cells in S phase, while endogenous Ki67 is expressed in cells throughout all cell cycle stages. As seen in prior work (Arguello et al., 2008; Lagace et al., 2010; Farioli-Vecchioli et al., 2012), this disconnect between these markers of proliferation may reflect a change in the number of cycling cells, or even the length of the cell cycle. Future work specifically assessing how mTBI changes SGZ cell cycle kinetics is warranted to address these possibilities.

The next time point, 7 dpi, was also selected due to it being consistently examined for TBI-induced changes in electrophysiology and behavior (Santhakumar et al., 2000, 2001; Schurman et al., 2017; Zhang et al., 2018). Indeed, the same LFPI parameters used here have been shown to result in DG hyperexcitability and deficits in DG-based cognition 7 dpi (Smith et al., 2012; Folweiler et al., 2018; Paterno et al., 2018), implying that hippocampal circuitry is disrupted in this mTBI model 7 dpi. An additional reason for choosing 7 dpi is that it allowed comparison to the progression of neurogenesis indices relative to 3 dpi. Based on the TBI-induced DG hyperactivity 7 dpi and the well-known ability of neuronal activity to stimulate neurogenesis (Kempermann, 2015; Kempermann et al., 2015; Káradóttir and Kuo, 2018), we expected to see more DCX+ cells 7 dpi, which we did **(Fig. 8)**. One interpretation of the increased DCX+ cells 7 dpi in the ipsilateral SGZ/GCL in LFPI vs. Sham mice is that at least some of the proliferating cells in the SGZ seen at 3 dpi express markers of immature neurons 4 days later at 7 dpi.

Previous work suggests that cells in the immature neuron stage at the time of injury (cells ‘born’ before the injury) are vulnerable to injury-induced cell death (Gao et al., 2008, 2009; Zhou et al., 2012; Hood et al., 2018). Our study did not directly assess the susceptibility of existing immature neurons to mTBI, as we were focused on the influence of mTBI on the process of neurogenesis after the injury. However, in this study, two pieces of our data suggest existing immature neurons are not susceptible to mild LFPI-induced cell death. First, if immature DCX+ cells were vulnerable to injury-induced death, we might expect to see a fewer DCX+ cells 3 dpi in the ipsilateral SGZ/GCL LFPI vs. Sham mice, as this population in part reflects cells born before the injury. However, we find DCX+ cell numbers are similar between Sham and LFPI mice 3 dpi. Second, if there was significant death of the DCX+ population after injury, we might expect a smaller GCL at the short-term time point; instead, we see similar DG GCL volume 3 dpi in Sham and LFPI mice. While these data suggest there is no gross loss of immature neurons due to the injury, future studies could directly address this by giving a BrdU injection prior to injury to label a cohort of cells born before the TBI, or by examining time points after the injury even earlier than 3 dpi.

In some models of injury, DCX+ cells have been reported outside the canonical DG neurogenic regions (the SGZ and inner GCL; Ibrahim et al., 2016; Shapiro, 2017). In our work here with an mTBI model, DCX+ cells in the oGCL were rare, thus we combined DCX+ cell counts in the inner and oGCL and SGZ. In addition, DCX+ cells in the hilus and molecular layer had ambiguous/faint staining, preventing reliable quantification of DCX+ cells in these non-canonical neurogenic DG regions. While this suggests our mTBI model leads to minimal DCX+ cells present outside of the SGZ and GCL, it is also possible that a more sensitive approach - such as a transgenic DCX+ reporter mouse - or distinct study design - such as labeling mitotic cells prior to injury - may reveal LFPI-induced DCX+ cells outside of these canonical DG neurogenic regions.

The final 31 dpi time point matches with evidence that the ipsilateral DG remains hyperexcitable 30 dpi (Tran et al., 2006) as well as the ~4 weeks it takes for adult-born granule cells to integrate fully into hippocampal circuitry (Brown et al., 2003; Gu et al., 2012). The BrdU injected 3 dpi intercalates into DNA of dividing cells such that staining for BrdU at 3 dpi will reveal the progeny of the initially-labelled cells that survived to 31 dpi. Here we report more BrdU+ cells in the ipsilateral SGZ 31 dpi in LFPI vs. Sham mice. As LFPI and Sham mice had the same number of BrdU+ cells 3 dpi **(Fig. 2 G-I)** and 7 dpi **(Fig. 4G-I)**, this emergence of more BrdU+ cells in LFPI vs. Sham SGZ 31 dpi combined with the overall fewer BrdU+ cells in 31 vs. 3 dpi tissue **(Fig. 6 G-I)** suggests an effect of survival **(Fig. 8)**. In other words, the rate of survival of BrdU+ cells in LFPI mice was higher (or the rate of death was lower) than the rate of BrdU+ in Sham mice. While the mechanisms mediating this difference are beyond the scope of this study, it is intriguing to consider if the TBI-induced hyperactivity reported 30 dpi (Tran et al., 2006) is responsible for this LFPI-induced survival, particularly given the well-described “activity-dependence” of adult-generated neurons (Deisseroth et al., 2004; Ma et al., 2009; Káradóttir and Kuo, 2018).

One caveat when discussing BrdU+ cells in the intermediate (7 dpi) and long-term (31 dpi) groups is that the DG subregion may influence the data interpretation. In the DG of naive adult male mice, BrdU has an *in vivo* half-life of 15 minutes (Mandyam et al., 2007), and thus BrdU labels cells that are in S-phase at the time of injection. In the SGZ, where the cell cycle is known to be 12 hours (Mandyam et al., 2007), we believe BrdU+ cells 7 and 31 dpi are post-mitotic, not cells that are still dividing. We favor this interpretation in the SGZ, as more than a few cell cycles would dilute the BrdU and make cells less likely to be detected via IHC (Dayer et al., 2003). Further supporting a post-mitotic state for SGZ BrdU+ cells 7 and 31 dpi is that our LFPI mice have more Ki67+ SGZ cells vs. Sham mice at 3 dpi, but LFPI and Sham Ki67+ SGZ cell numbers are similar at 7 and 31 dpi. Beyond being post-mitotic, we also suspect BrdU+ SGZ cells 31 dpi are mature granule cell neurons, but this speculation merits more discussion (see the first ‘future direction’ below). Outside of the SGZ, it is more likely that BrdU injection 3 dpi labels proliferating cells whose progeny become non-neuronal cells. Certainly, reactive gliosis is evident in many brain regions in other and more severe TBI models (Acosta et al., 2013; Newell et al., 2019; Caplan et al., 2020). The spatiotemporal extent and severity of reactive astro- and micro-gliosis in DG subregions in this mild LFPI model is less well-studied, although there is reactive astrocytosis in the ipsilateral hippocampus 7 dpi (Smith et al. 2015). Therefore, it is possible (and even likely) that dividing glia or other brain cells take up BrdU (from the BrdU injection administered 3 dpi) and persist in the hilus, oGCL, and molecular layer, DG subregions which have distinct cell cycle kinetics from the SGZ (Mandyam et al., 2007), and thus whose progeny may still be proliferating 7 and 31 dpi. If these progeny of hilus, oGCL, and molecular layer cells do become post-mitotic, it is also possible their non-neuronal post-mitotic progeny survive to 7 and 31 dpi. While these are reasonable speculations, these possible DG subregion-dependent state and fate outcomes warrant testing with cell cycle markers in both Sham and LFPI tissue, and usually multiple time points. Minimally, though, our data suggest the importance of considering the dynamics and DG subregion-dependent effects of injury-induced neurogenesis.

While we report LFPI leads to transient, sequential changes in proliferating cells, neuroblasts/immature neurons, and surviving cells, we find no LFPI-induced change in DG granule cell layer volume **(Table 1)**. Volume is an important consideration since more new cells may be compensating for TBI-induced cell loss caused by the injury. However, our data suggest there is no decrease in the size of the GCL, and therefore no major tissue loss, raising the possibility of overcompensation by the brain. Additionally, many papers examining neurogenesis after TBI report neurogenic cell counts in terms of density, not in terms of stereologically-determined cell number. While density is a useful metric, reporting both GCL volume and cell counts allows for comparison with those studies while preserving data resolution (increased density of new neurons could be from either increased number of cells or a decrease in volume). Therefore, our data demonstrating more neurogenesis in the absence of increased GCL volume reflects a true increase in the neurogenic rate, rather than an overall increase in DG processes or capacity.

Our findings raise several important future directions. First, it will be important to definitively determine how mTBI influences the fate and integration of adult-generated DG granule cells into DG circuitry. As for the fate of these cells, there are several reasons we interpret the LFPI-induced increase in BrdU+ SGZ cells 31 dpi as reflecting increased survival specifically of adult-born DG granule cells. This interpretation is supported by the size, shape, and location of the BrdU+ cells. The BrdU+ nuclei are the right size (~10um) to be neuronal nuclei of DG granule neurons. The cell bodies of the BrdU+ nuclei are circular, suggestive of neurons (or at least not suggestive of reactive astrocytes or other glia), and they are in the SGZ and inner GCL where most adult-generated neurons end up (Kempermann et al., 2003). While it is possible that some of the BrdU+ cells 31 dpi outside of the neurogenic SGZ/GCL are glia or other brain cells, most astrocytes and microglia reside in the hilus and molecular layer and only sparsely populate the GCL. Nevertheless, conclusive determination of the neuronal fate of BrdU+ SGZ cells 31 dpi will require phenotypic analysis of BrdU+ SGZ cells 31 dpi with a neuronal marker and/or electrophysiological characterization. Second, as other models of TBI and seizure models result in aberrant migration of adult-born DG neurons (Dashtipour et al., 2003; Emery et al., 2005; Shapiro et al., 2008, 2011; Murphy et al., 2012; Ibrahim et al., 2016; Ratliff et al., 2020), future work could assess whether the greater number of BrdU+ cells in DG subregions adjacent to the SGZ (hilus and oGCL) means this may also be occurring in our mTBI model. In our study, DCX+ staining in non-neurogenic regions was ambiguous, but future work using transgenic or viral-mediated visualization of DCX+ cells or the neuronal phenotyping mentioned above would be useful in this regard (van Praag et al., 2002; Laplagne et al., 2006; Wood et al., 2011; Kathner-Schaffert et al., 2019; Cole et al., 2020; Tensaouti et al., 2020). Third, it will be important to assess if our mTBI model changes the dendritic morphology of the adult-born DG granule cells as has been reported in other and more severe TBI models (Folkerts et al., 1998; Gao et al., 2011; Villasana et al., 2015; Ibrahim et al., 2016). In this mTBI model, there were no injury-induced gross visual differences in DCX+ process length, thickness, or angle. However, our stained DG tissue was marked by a high density of immature neuron processes, typical of DCX+ staining in young adult mice (Wu and Hen, 2014; Villasana et al., 2019), and this prevented performing a rigorous quantitative analysis of dendritic morphology. Future studies could more carefully address the impact of mTBI on dendritic morphology using alternative methods (i.e., fluorescent retroviral labeling) to more sparsely label new DG neurons, allowing for high-resolution analysis of newborn cell dendritic morphology.

A final future direction will be to clarify the question of functional significance of the injury-induced transient surge of proliferating cells, immature neurons, and surviving cells shown here. Certainly, in uninjured animals, DG neurogenesis is critical for cognitive function (Clelland et al., 2009; Sahay et al., 2011). After injury, the prevailing idea is that increased neurogenesis represents an attempt by the brain to repair itself, and thus it aids functional recovery (Blaiss et al., 2011; Sun et al., 2015). However, preclinical work in seizures (Cho et al., 2015) and TBI (Neuberger et al., 2017) have questioned the assumption that more new neurons are always beneficial, and underscore that the timing and methods of manipulating new DG neurons may influence the outcome of such studies. Because adult-born neurons are exquisitely sensitive to external cues (Ming and Song, 2005; Aimone et al., 2014; Bond et al., 2015; Kempermann et al., 2015; Catavero et al., 2018), the pathological environment of a post-injury dentate has the potential to influence newborn neuron development in a maladaptive manner. Therefore, the question of reparative potential of injury-induced neurogenesis needs to be explicitly tested in a less invasive manner than ablation of new neurons; chemogenetic or optogenetic manipulation of injury-induced neurons would allow more accurate inquiry into how injury-induced neurons influence hippocampal function.

Together, these data suggest this model of mTBI induces transient, sequential increases in ipsilateral SGZ/GCL proliferating cells, immature neurons, and surviving cells which we interpret as a transient increase in mTBI-induced neurogenesis. Our work in this mTBI model is in line with other and more severe models of TBI, where the large fraction of literature shows injury-induced stimulation of DG neurogenesis. As our data in this mTBI model provide temporal, subregional, and neurogenesis-stage resolution, these data are important to consider in regard to the functional importance of TBI-induced neurogenesis and future work assessing the potential of replacing and/or repairing DG neurons in the brain after TBI. Indeed, the transient nature of the injury-induced increase in neurogenesis provides temporal guidance about a discrete window for targeting injury-induced adult-born neurons. Because DCX+ cells are increased 7 dpi, but not 3 or 31 dpi, it is the neurons born during that intermediate timeframe that have the most potential to disrupt the circuit. This is good news for future work attempting to target these cells for chemo- or optogenetic targeting, as it narrows down the temporal window and population subset for targeting. We hope future work will consider the “age” of the cells, time post-injury, and anatomical location and integration of the cells, as these parameters may prove useful to achieve therapeutic potential of any intervention strategy.

---

**Figure S1.**
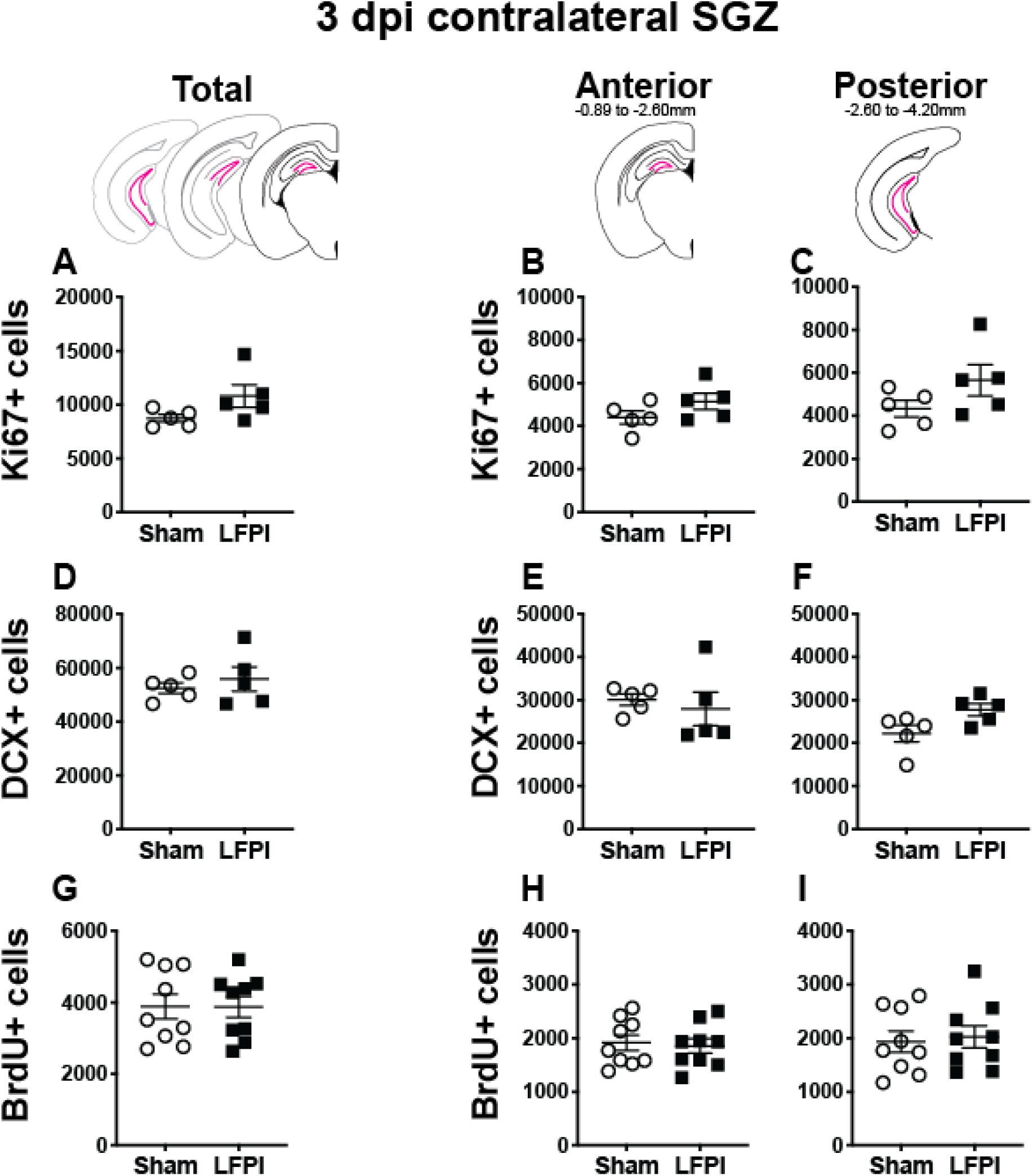
Relative to Sham, lateral fluid percussion injury (LFPI) does not change the number of Ki67-immunoreactive (Ki67+), DCX+, or BrdU+ cells in the contralateral mouse subgranular zone (SGZ) or granule cell layer (GCL) 3 days post-injury (dpi). Pink lines in schematics (top row) indicate these measures were collected in the contralateral SGZ/GCL, while the green lines in the figures in the main text indicate those measures were collected in the ipsilateral SGZ/GCL. Stereological quantification of Ki67+ **(A-C**; Sham n=5, LFPI n=5), DCX+ **(D-F**; Sham n=5, LFPI n=5), and BrdU+ **(G-I**; Sham n=9, LFPI n=9)cells in the SGZ (Ki67, BrdU) and GCL (DCX). Immunopositive cells were quantified across the entire longitudinal axis **(A, D, G)**, and also broken up into anterior **(B, E, H)** and posterior **(C, F, I)** bins, operationally defined as Bregma levels −0.92 to −2.6; and −2.6 to −3.97, respectively.

**Figure S2.**
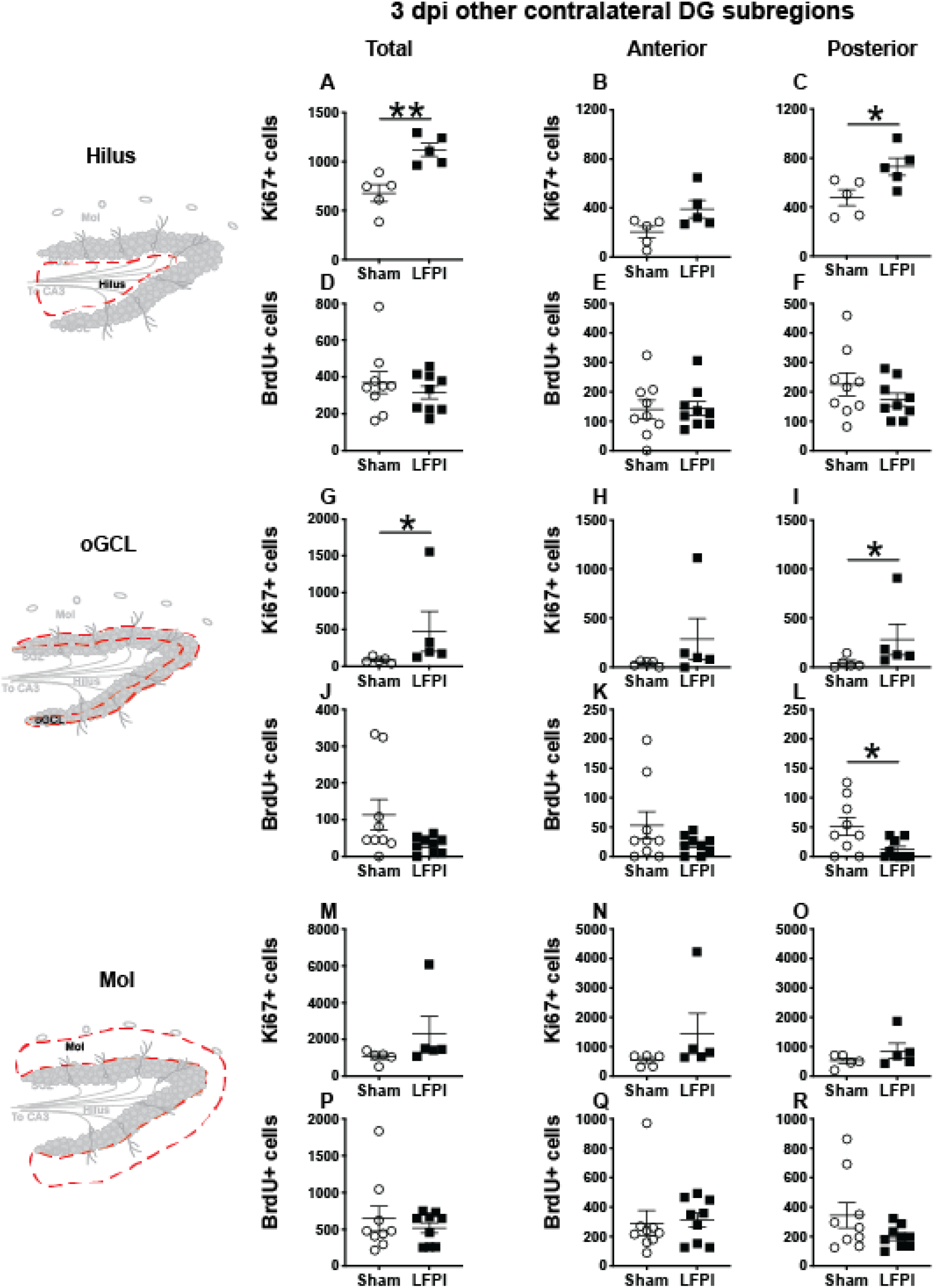
Relative to Sham, LFPI increases proliferation in some contralateral other DG subregions 3 days post-injury. Stereological quantification of Ki67+ **(A-C, G-I, M-O**; Sham n=5, LFPI n=5)and BrdU+ **(D-F, J-L, P-R**; Sham n=9, LFPI n=9)cells in the hilus (**A-F**; red dotted line region, top-left schematic), outer granule cell layer (**G-L**; red dotted line region, middle-left schematic), and molecular layer (**M-R**; red dotted line region, bottom-left schematic). Immunopositive cells were quantified across the entire longitudinal axis **(A, D, G, J, M, P)**, and also broken up into anterior **(B, E, H, K, N, Q)** and posterior **(C, F, I, L, O, R)** bins, operationally defined as Bregma levels −0.92 to −2.6; and −2.6 to −3.97, respectively. One way ANOVA, *p<0.05, **p<0.01.

**Figure S3.**
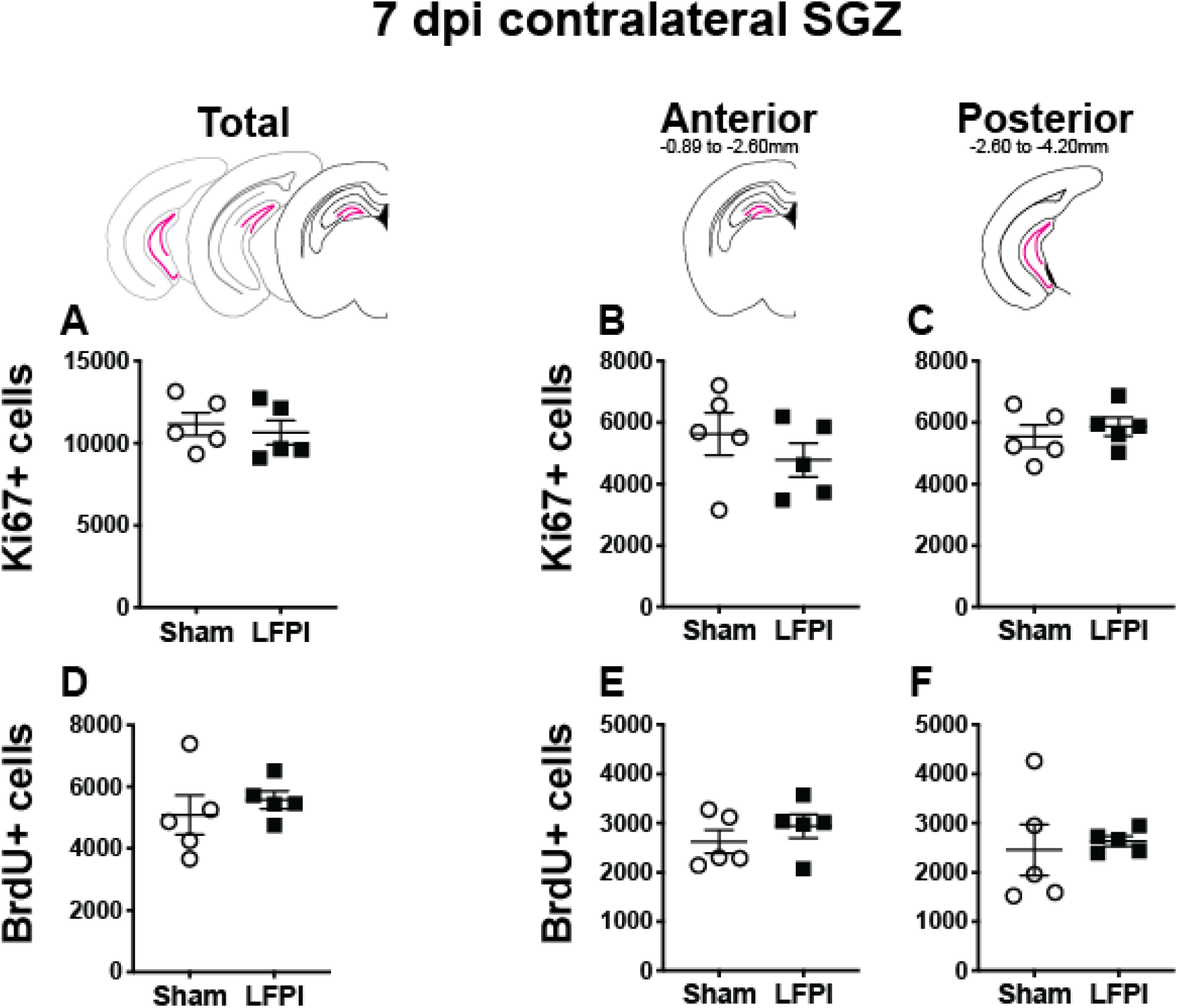
LFPI does not affect proliferation and cell survival in the contralateral mouse SGZ 7 days post-injury. Pink lines in schematics (top row) indicate these measures were collected in the contralateral SGZ/GCL. Stereological quantification of Ki67+ **(A-C**; Sham n=5, LFPI n=5)and BrdU+ **(D-F**; Sham n=5, LFPI n=5)cells in the SGZ (Ki67, BrdU) and GCL (DCX). Immunopositive cells were quantified across the entire longitudinal axis **(A, D)**, and also broken up into anterior **(B, E)** and posterior **(C, F)** bins, operationally defined as Bregma levels −0.92 to −2.6; and −2.6 to −3.97, respectively.

**Figure S4.**
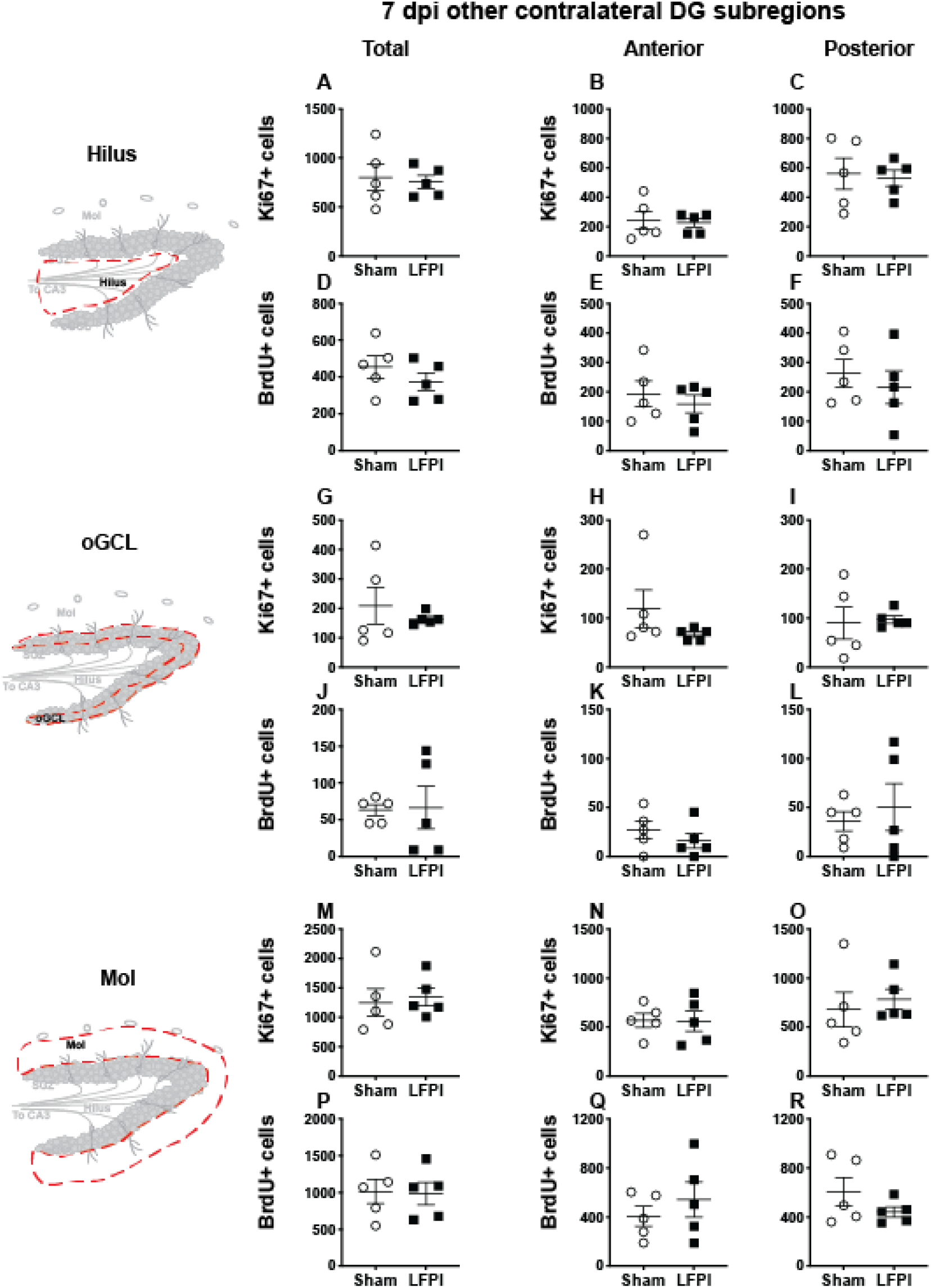
LFPI does not affect proliferation and intermediate cell survival in other DG subregions 7 days post-injury. Stereological quantification of Ki67+ **(A-C, G-I, M-O**; Sham n=5, LFPI n=5)and BrdU+ **(D-F, J-L, P-R**; Sham n=5, LFPI n=5)cells in the hilus (**A-F**; red dotted line region, top-left schematic), outer granule cell layer (**G-L**; red dotted line region, middle-left schematic), and molecular layer (**M-R**; red dotted line region, bottom-left schematic). Immunopositive cells were quantified across the entire longitudinal axis **(A, D, G, J, M, P)**, and also broken up into anterior **(B, E, H, K, N, Q)** and posterior **(C, F, I, L, O, R)** bins, operationally defined as Bregma levels −0.92 to −2.6; and −2.6 to −3.97, respectively.

**Figure S5.**
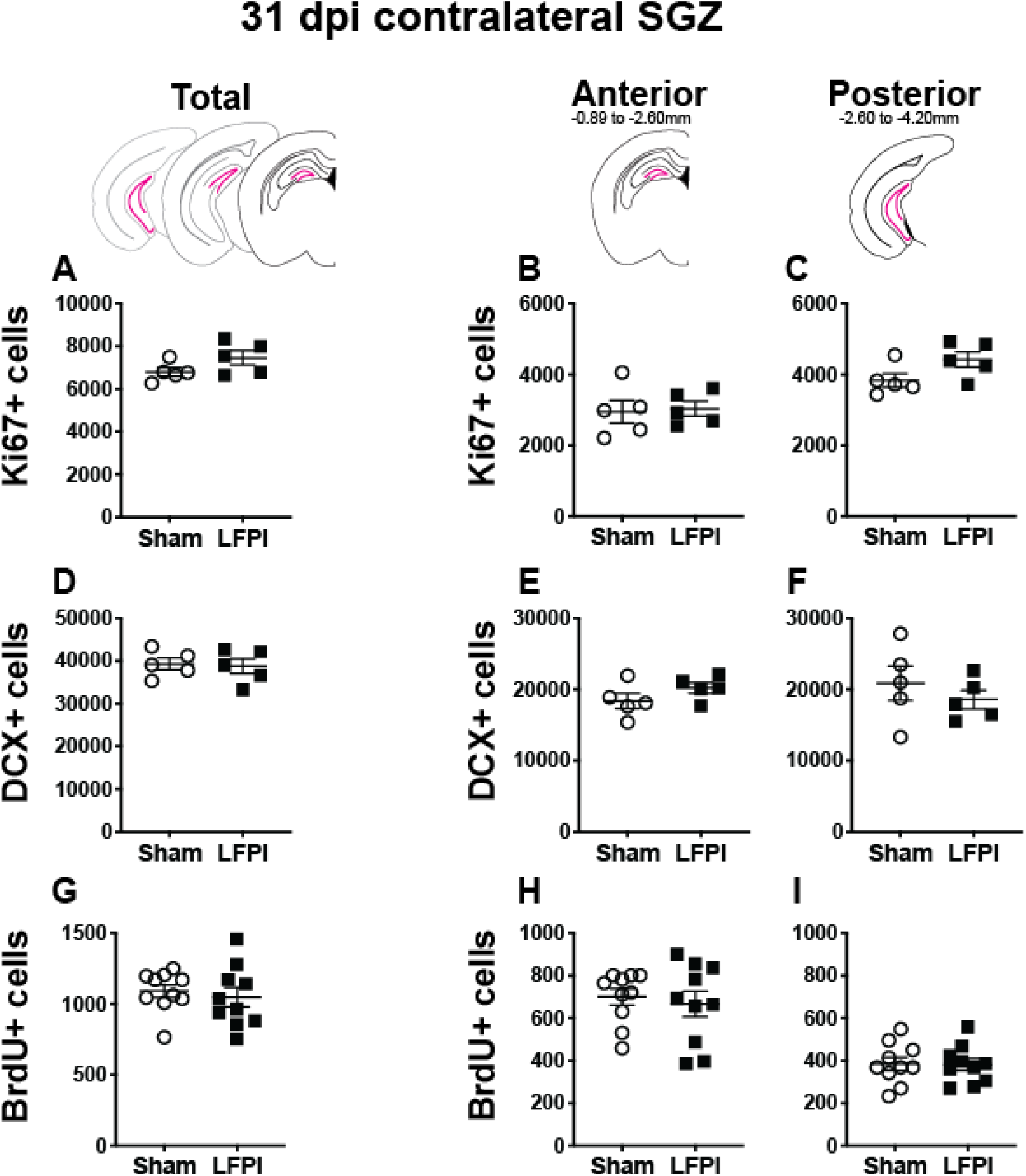
LFPI does not affect proliferation, neurogenesis, or long-term cell survival in the contralateral mouse SGZ 31 days post-injury. Pink lines in schematics (top row) indicate these measures were collected in the contralateral SGZ/GCL. Stereological quantification of Ki67+ **(A-C**; Sham n=5, LFPI n=5), DCX+ **(D-F**; Sham n=5, LFPI n=5), and BrdU+ **(G-I**; Sham n=10, LFPI n=10)cells in the SGZ (Ki67, BrdU) and GCL (DCX). Immunopositive cells were quantified across the entire longitudinal axis **(A, D, G)**, and also broken up into anterior **(B, E, H)** and posterior **(C, F, I)** bins, operationally defined as Bregma levels −0.92 to −2.6; and −2.6 to −3.97, respectively.

**Figure S6.**
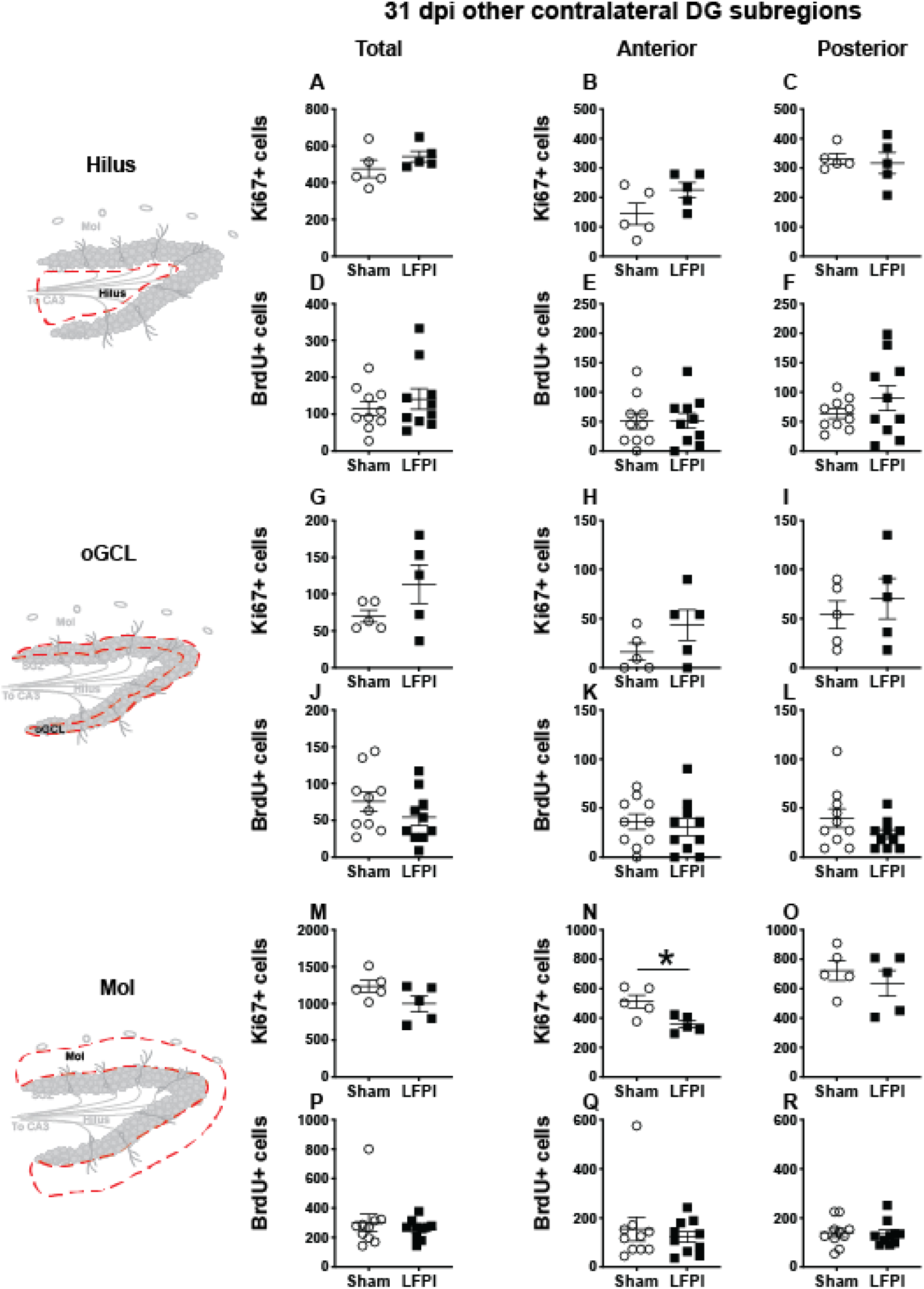
LFPI does not affect proliferation or long-term cell survival in other contralateral DG subregions 31 days post-injury. Stereological quantification of Ki67+ **(A-C, G-I, M-O**; Sham n=5, LFPI n=5)and BrdU+ **(D-F, J-L, P-R**; Sham n=10, LFPI n=10)cells in the hilus (**A-F**; red dotted line region, top-left schematic), outer granule cell layer (**G-L**; red dotted line region, middle-left schematic), and molecular layer (**M-R**; red dotted line region, bottom-left schematic). Immunopositive cells were quantified across the entire longitudinal axis **(A, D, G, J, M, P)**, and also broken up into anterior **(B, E, H, K, N, Q)** and posterior **(C, F, I, L, O, R)** bins, operationally defined as Bregma levels −0.92 to −2.6; and −2.6 to −3.97, respectively.

**Supplementary Table 1.**
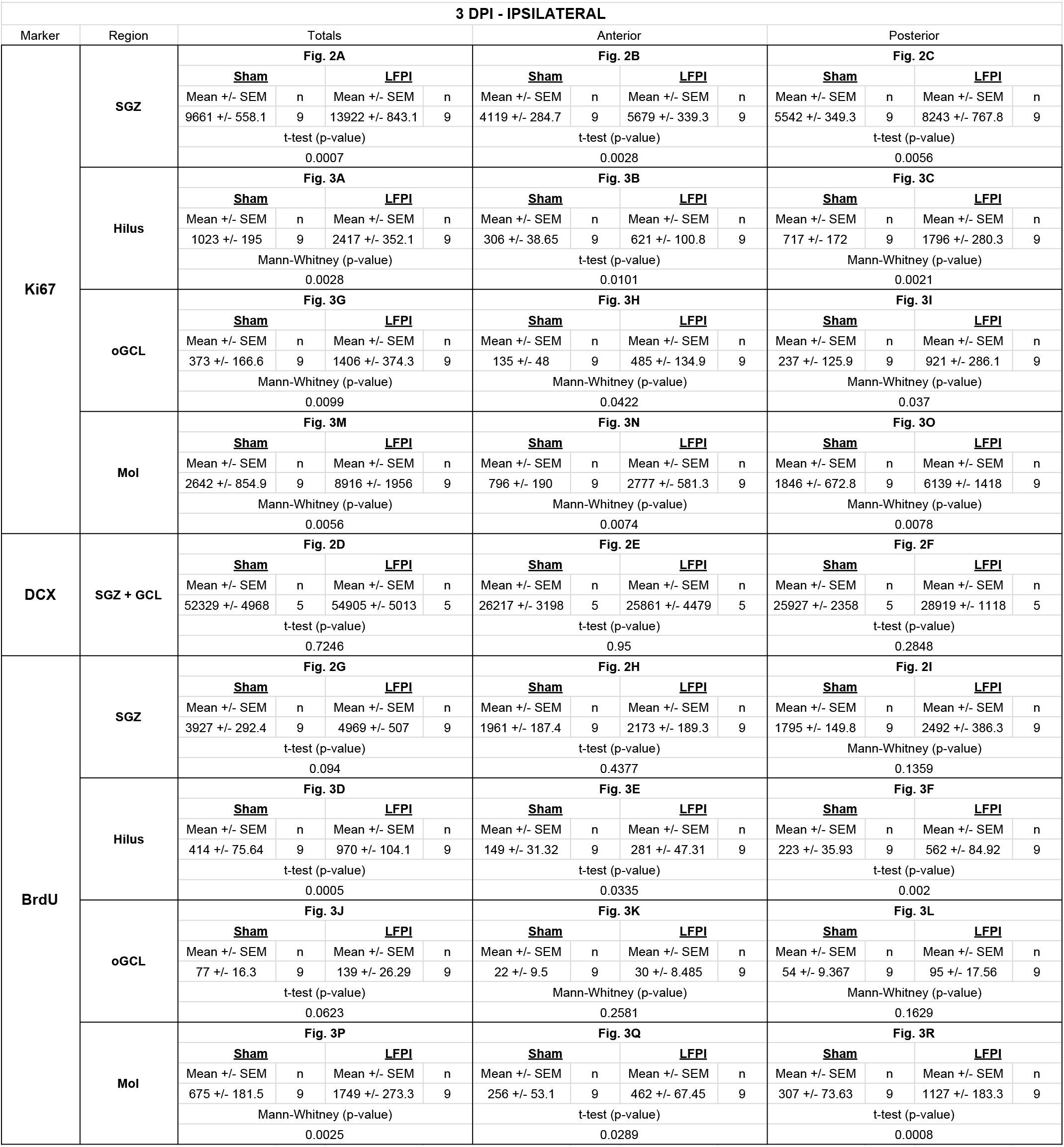
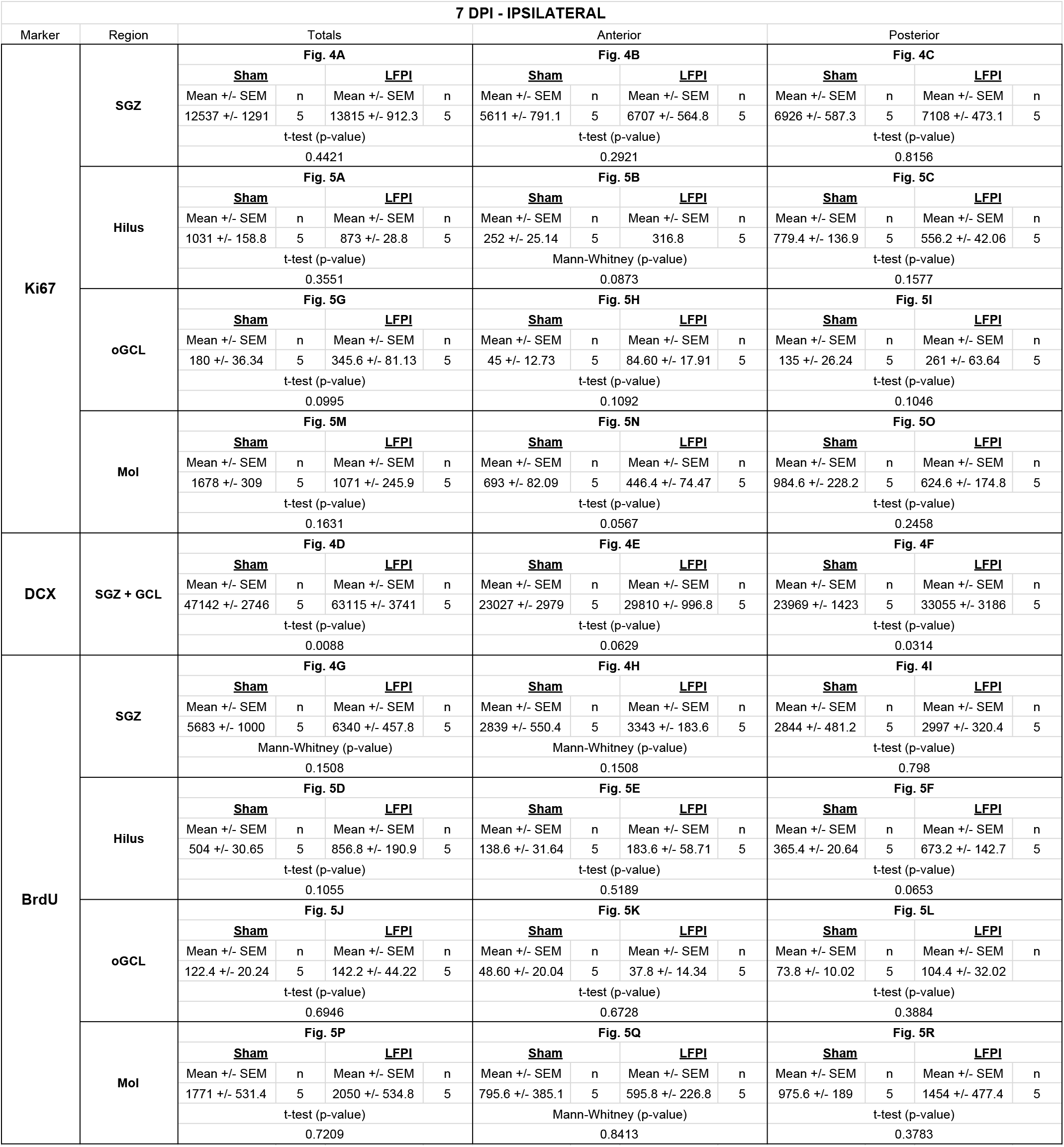
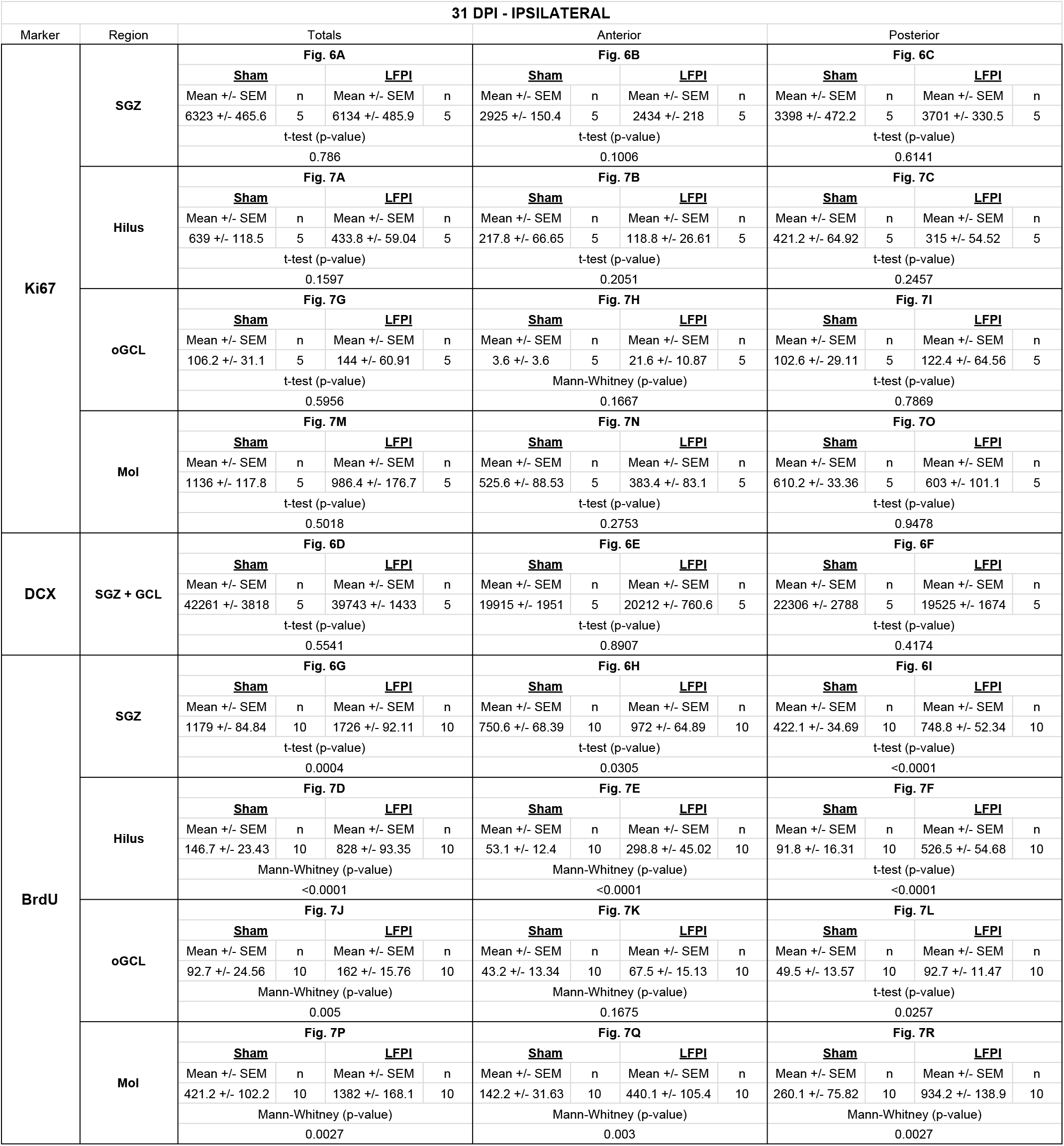
Statistical results for cell quantification in the ipsilateral hemisphere.

**Supplementary Table 2.**
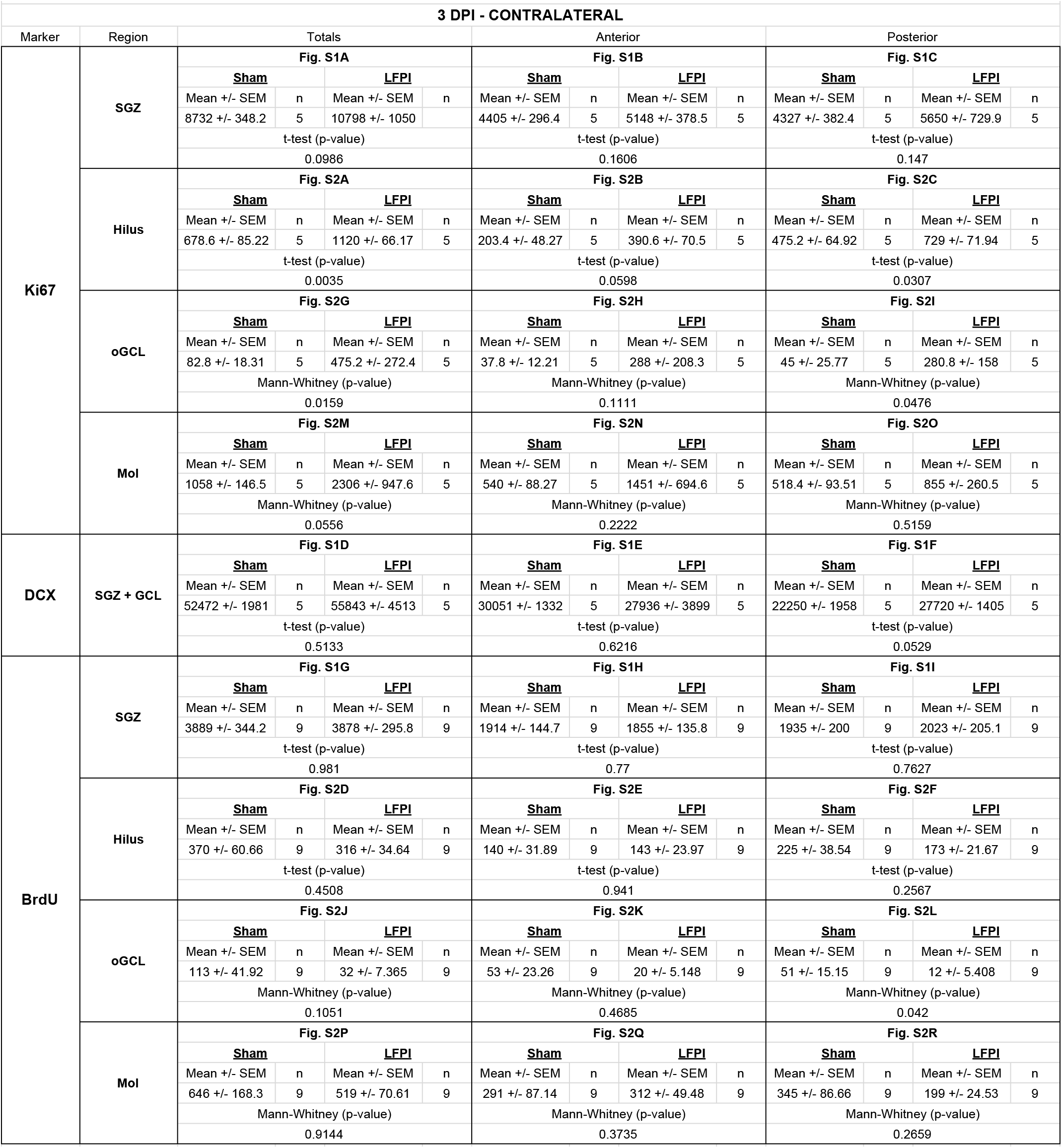
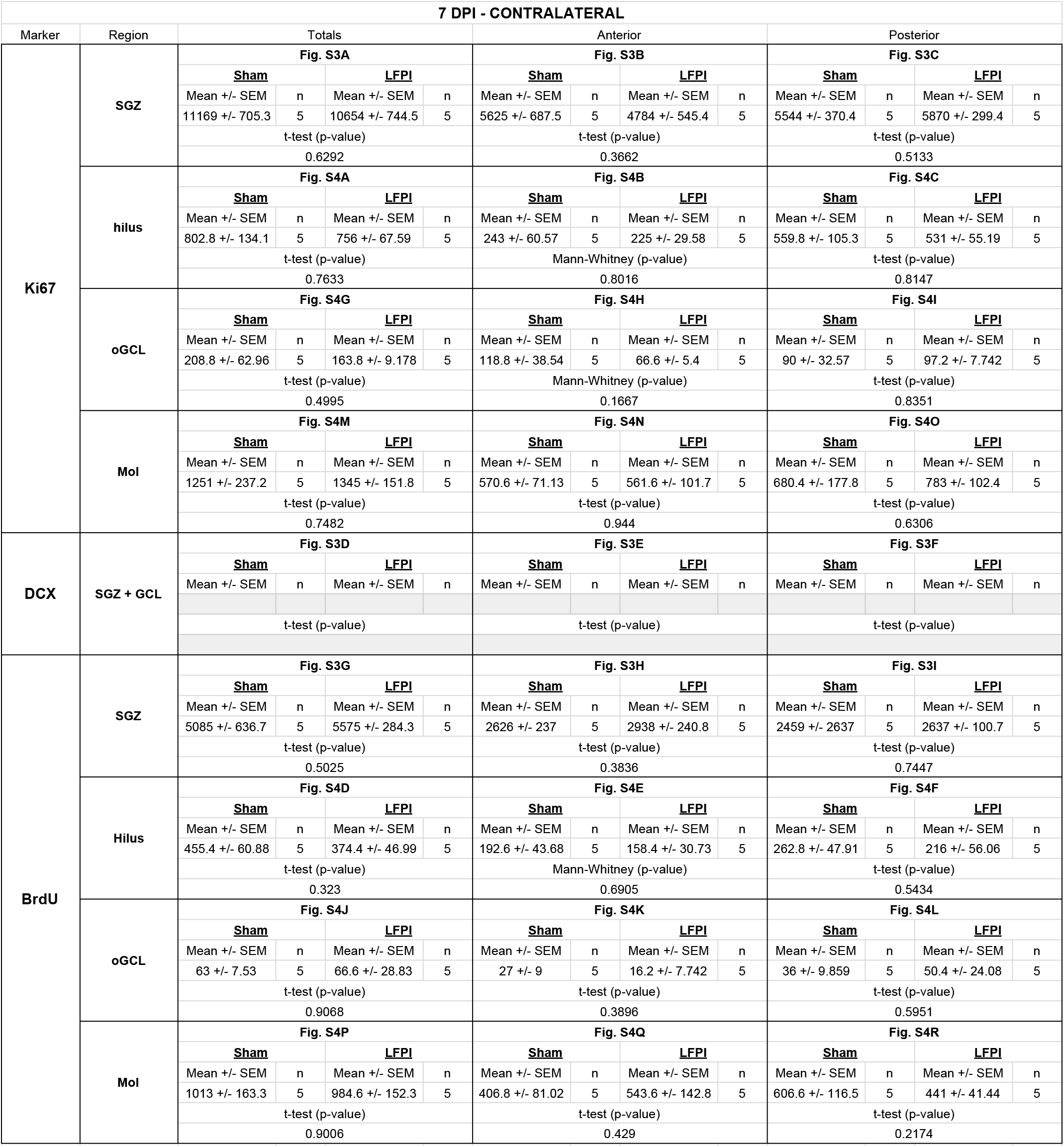
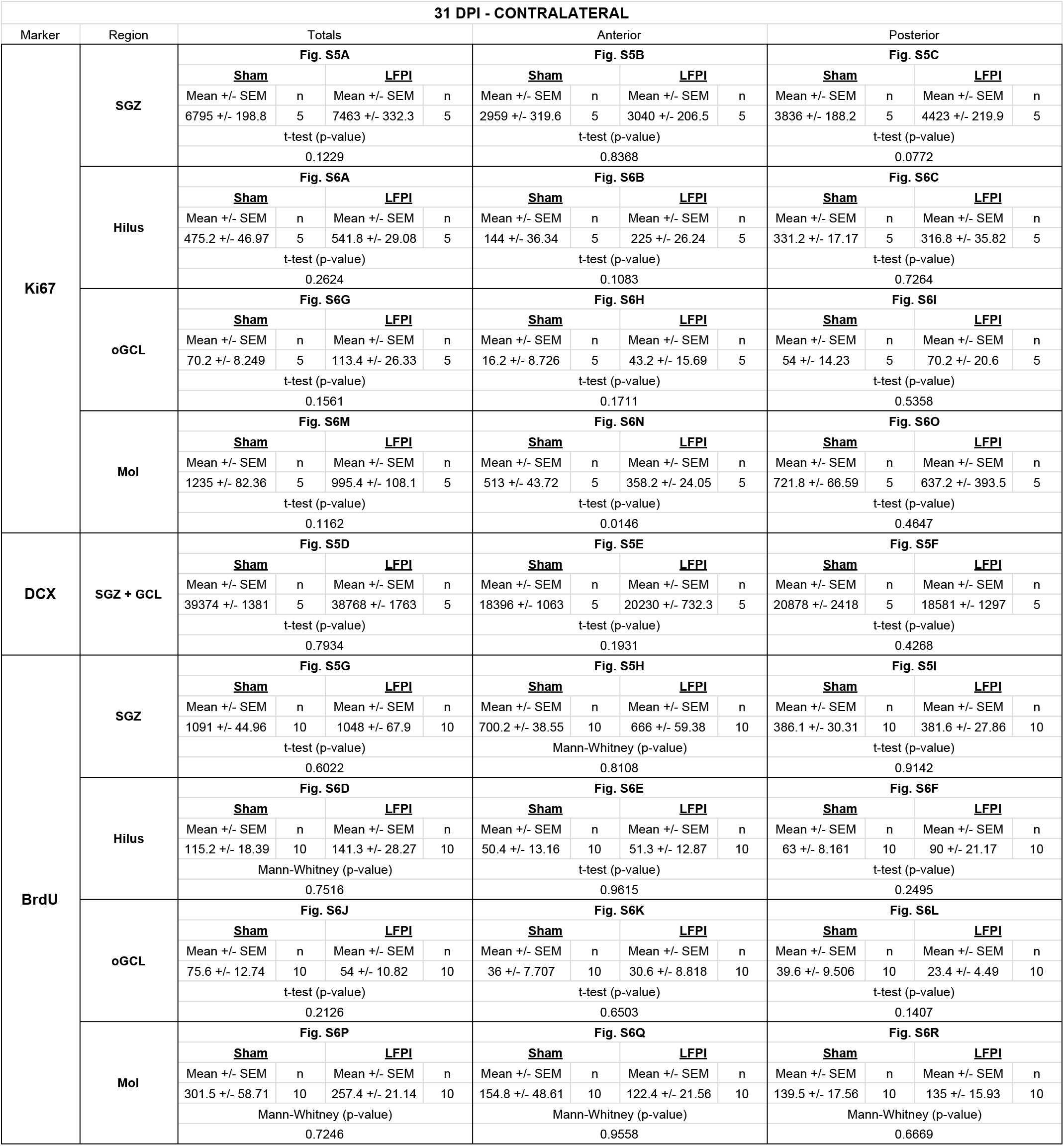
Statistical results for cell quantification in the contralateral hemisphere.

